# Common fragile sites are characterised by faulty condensin loading after replication stress

**DOI:** 10.1101/508713

**Authors:** Lora Boteva, Ryu-Suke Nozawa, Catherine Naughton, Kumiko Samejima, William C Earnshaw, Nick Gilbert

## Abstract

Cells coordinate interphase to mitosis transition but recurrent cytogenetic lesions appear at common fragile sites (CFSs) in a tissue-specific manner following replication stress, marking regions of instability in cancer. Despite such a distinct defect no model fully explains their molecular configuration. We show that CFSs are characterised by impaired chromatin folding manifested as disrupted mitotic structures visible using molecular FISH probes in the presence and absence of replication stress. Chromosome condensation assays reveal that compaction-resistant chromatin lesions persist at CFSs throughout the cell cycle and mitosis. Subsequently cytogenetic and molecular lesions arise due to faulty condensin loading at CFSs, through a defect in condensin I mediated compaction and are coincident with mitotic DNA synthesis (MIDAS). This model suggests that in conditions of exogenous replication stress, aberrant condensin loading leads to molecular defects and CFS formation, concomitantly providing an environment for MIDAS, which, if not resolved, result in chromosome instability.

## Introduction

The folding of chromosomes in preparation for mitosis is the most profound structural change the genome undergoes throughout a cell’s lifetime (Antonin and Neumann, 2016). Mitotic condensation is linked to successful cell division and cell cycle progression in a functional and regulatory manner and its failure can be costly, leading to lagging chromosomes and aneuploidy (Gordon et al., 2012; Zhu et al., 2018). Much effort has been made to define the molecular basis of the condensation process and bridge the cytogenetic features of mitotic chromosomes with molecular-level understanding of the chromatin and scaffolding proteins that comprise them. As a result, it is now accepted that a fully folded, cytogenetically normal metaphase chromosome is the product of successful and timely completion of inter-connected processes including replication, sister chromatid separation and chromatin condensation (Wechsler et al., 2011; Ono et al., 2013; Gibcus et al., 2018). Consequently, the cytogenetic integrity of chromosomes is affected by the disruption of these processes; common fragile sites (CFSs)(Durkin and Glover, 2007), regions of the genome known for forming lesions on metaphase chromosomes when cells are challenged with replication stress (Zeman and Cimprich, 2014), are a prominent example. Illustrating the importance of mitotic compaction for genome stability, these sites overlap with recurrent cancer deletions and tumour suppressor genes frequently lost in cancer (Le Tallec et al., 2013; Bignell et al., 2010; Negrini et al., 2010). Unlike constitutively fragile locations such as Fragile X, CFSs form in a cell type specific manner, leading to suggestions that an epigenetic component plays a role in their fragility (Letessier et al., 2011; Le Tallec et al., 2011). Although, the mechanistic basis for their fragility is still unknown a number of shared factors have been identified, including late replication timing, transcription of long genes and features of the underlying DNA sequence (Helmrich et al., 2011; Le Tallec et al., 2014; Fungtammasan et al., 2012; Wilson et al., 2015; Blin et al., 2019; Wei et al., 2016).

CFSs require FANCD2 for efficient replication (Madireddy et al., 2016) and have been identified as regions where active DNA synthesis is apparent on mitotic chromosomes in a process dependent on POLD3 and the Mus81 nuclease (Minocherhomji et al., 2015). The exact steps involved in triggering synthesis remain unknown but it also requires the TRAIP ubiquitin ligase (Sonneville et al., 2019) and may be active at regions of the genome that are incompletely condensed in mitosis. Recently, condensation defects have also been shown to underlie HR-deficiency mediated mitotic lesions, and if not resolved lead to DNA damage and chromosomal instability (Chan et al., 2018). The effectors of such condensation failures are likely to be proteins that drive mitotic folding such as the condensin I and II complexes which are crucial for chromosome compaction (Gibcus et al., 2018; Samejima et al., 2012; Lipp et al., 2007). Furthermore, mechanisms established in yeast show that condensin loading is inhibited in slow replicating zones, which resemble CFS, by the ATR homologue Mec1 (Cha and Kleckner, 2002; Hashash et al., 2012).

Given the close relationship between replication and mitotic compaction (Ono et al., 2013), we hypothesised that disrupted mitotic folding may arise as a consequence of replication stress at sensitive regions such as CFS. Using a FISH-based approach, we show that CFS are characterised by failure of local chromatin to compact for mitosis: this is the case at cytogenetic lesions but also when the sites appear cytogenetically normal, where we demonstrate a previously unknown propensity for smaller scale molecular lesions (100 kb), visible only at the molecular (imaged by FISH) and not the cytogenetic level. We show that molecular and cytogenetic instability at CFSs is mechanistically dependent on a failure to recruit condensin and remodel chromatin at the G2/M boundary to facilitate mitotic folding. Analysis of specific condensin complexes indicates that condensin I, rather than condensin II, is the effector of disrupted mitotic compaction at CFSs. Our model suggests that after replication non-fragile regions undergo structural and compositional ‘priming’ of chromatin in preparation for mitosis. In contrast CFSs are regions of the genome where, even in unperturbed conditions, chromatin is inefficiently ‘primed’ for mitotic compaction, characterised by aberrant condensin loading, leading to molecular lesions, whilst in the extreme conditions of exogenous replication stress cytological chromosome abnormalities are apparent.

## Results

### CFS frequency and repertoire in RPE1 and HCT116 cells

To analyse the relationship between chromosome architecture and CFS structure we characterised the CFS repertoire and frequency in two epithelial chromosomally near-normal diploid cell lines (HCT116 and RPE1), using DAPI banding, after inducing replication stress with aphidicolin (APH). 372 lesions across 371 metaphases for APH concentrations ranging from 0.1 to 0.6 μM were observed, showing that higher APH concentration led to increased rate of breakage and more severe CFS phenotypes (Supplementary Figure 1A-B) but did not cause cell cycle arrest (Supplementary Figure 1C). Cytogenetic lesions were then mapped and scored in metaphase spreads prepared from HCT116 (n= 94) and RPE1 (n = 64) cells following 24-hour treatment with 0.4 μM APH (Figure 1A-B, Supplementary Figure 1D-E, Supplementary Table 1). Despite both cell lines being of epithelial origin, the CFS repertoire differed significantly: FRA3B was the most fragile site in the HCT116 line (23% of all breaks), followed by locations on chromosome 2 (FRA2I, 2q33.2; FRA2T, 2q24.1). In contrast, the most fragile location in the RPE1 cell line, FRA1C on 1p31.2 was only weakly fragile in HCT116 (18.6% of all breaks in RPE1; 5.8% in HCT116); additionally, one of the most common break sites (approx. 10% of all breaks) in the RPE1 cell type have not been previously identified as a CFS location: 4q32.2. A previous analysis of CFS distribution in HCT116 cells (Le Tallec et al., 2013) also indicated that FRA3B was the most common site but there were also differences: in our study FRA4F and FRA2I instability were more frequent whilst FRA4D and FRA16D instability was not readily apparent. In contrast, a further study found that FRA16D was the most common fragile site in HCT116 cells (Hosseini et al., 2013) indicating differences in CFS repertoire and frequency between sub-clones.

CFSs are reported to share a number of structural characteristics: the presence of long genes, AT-rich sequences and late replication timing (Fungtammasan et al., 2012; Arlt et al., 2009; Wilson et al., 2015). The genomic features of the sites we identified were consistent with these trends, although CFSs do not contain either the most GC-poor regions in the genome or the longest genes (Figure 1C-D). Among the most fragile locations in our study, 9 out of 11 overlapped with genes larger than 0.3 Mb including FRA3B (FHIT), FRA4F (GRID2, CCSER), 4q32.2 (MARCH1, 0.85 Mb; FSTL5, 0.78 Mb) and FRA7E, which spans MAGI2 (1.4 Mb). While the frequent FRA1C site in the RPE1 cell line does not overlap with any long genes, it is in close proximity to LRRC7 (0.32 Mb) and 2.5 Mb away from NEGR1 (0.89 Mb). COSMIC mutation data also showed that, as expected, the majority of the most frequent CFSs (8 out of 11) overlapped with recurrent cancer deletion clusters (Le Tallec et al., 2013).

**Figure 1:**
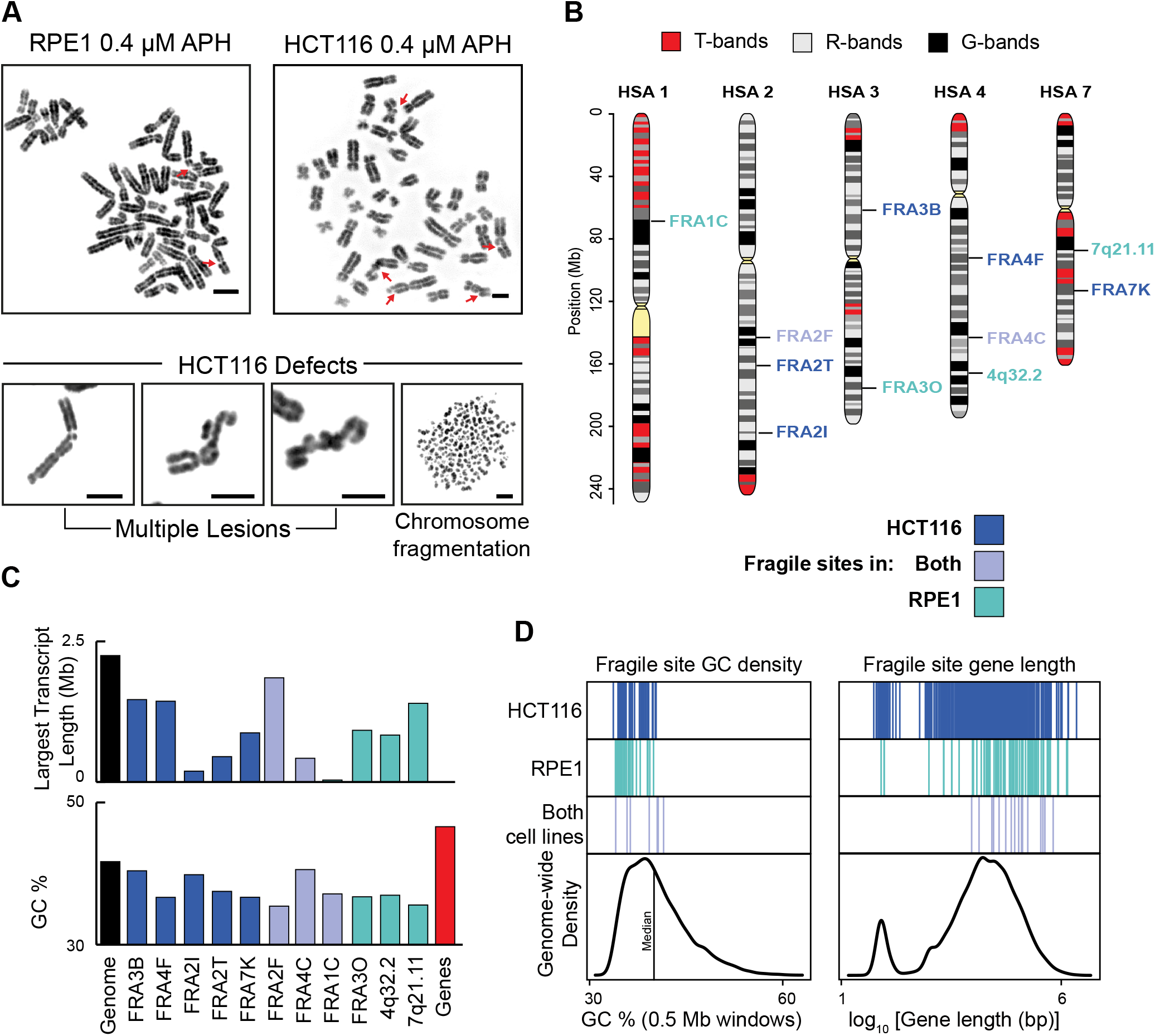
Characterisation of CFSs in HCT116 and RPE1 epithelial cells. **A.** Ideograms showing most frequent aphidicolin dependent common fragile site locations in RPE1 and HCT116 epithelial cells, cytogenetically scored using DAPI banding. CFSs specific to HCT116 cells (blue), RPE1 (green) and both (mauve) are indicated. **B**. Representative metaphase spreads (reverse DAPI banding) from RPE1 (left) and HCT116 (right) cell lines, showing CFS fragility (red arrows) following aphidicolin treatment (top); bottom, extreme chromosomal defects in HCT116 cells; Scale bar, 5 μm. **C**. Length of largest transcript (top) and GC content (bottom) at sites fragile in HCT116 (blue), RPE1 (green) or both cell lines (mauve). Scale bar, 5 μm. **D**. Left, genome-wide GC (% in 0.5 Mb windows) density plot with GC density at CFSs in HCT116 (blue), RPE1 (green) or both cell lines (mauve). Right, genome-wide gene length (NCBI genes) density plot with gene length of genes encompassed within CFSs in HCT116 (blue), RPE1 (green) or both cell lines (mauve).

### CFS regions have irregular chromatin structures in the absence of replication stress

Cytogenetic mapping after replication stress revealed a range of lesion phenotypes at mitosis including chromatid breaks and gaps, chromosome gaps, concatenations and other complex abnormalities, with no relationship between particular locations and the type of abnormality observed (Figure 1). As cytogenetic mapping provides relatively low-resolution information on the molecular location of a fragile site lesion a BAC-walking strategy was used to fine-map five cytogenetically identified CFS regions (Supplementary Table 1). Probes were selected spanning the sites and the frequency of chromosomes showing cytogenetic lesions overlapping with the probes were quantified (Figure 2A, Supplementary Figure 2A). Rather than always occurring at the same location, breaks appeared across large genomic regions encompassing CFS sites: a high frequency of breaks were observed at a fragile “core” region, which tailed off at BACs located upstream or downstream (80% break overlap at the core of the sites reduced to 33% at the flanks). Fluorescent BAC signals were often observed to span CFS lesions, with the fluorescence intensity of the probes peaking over the DAPI faint regions, consistent with DNA being present within the cytogenetically visible breaks (Supplementary 2B).

To better characterise mitotic chromosome fragility the chromatin state of CFSs was assessed using the FISH signal from the BAC probes as a marker for chromatin condensation, at a molecular level. After aphidicolin treatment probes within CFS regions showed a propensity to have atypical FISH signals even in the absence of cytogenetic lesions at the corresponding CFS site, (Figure 2B and Supplementary Figure 2D). Rather than twin symmetric foci on mitotic chromosomes, CFS spanning probes frequently formed multiple, asymmetric spots, or appeared as a single spot sitting between the two chromatids. However, the most extreme of these atypical signals was a phenotype in which BACs extended away from the chromosome, spreading far beyond the DAPI-dense area. Although aberrant chromatin folding is often seen in FISH of mitotic chromosomes under the conditions used for these studies control loci rarely showed chromatin compaction defects (Supplementary Figure 2D). Furthermore, these irregular structures are unlikely to be a consequence of using a specific probe as aberrant FISH signals were observed clustered across fragile sites. Instead these signals are reminiscent of abnormal FISH signals formed at telomeres in response to replication stress termed “fragile telomeres” (Sfeir et al., 2009) and are indicative of problems with mitotic condensation and decatenation.

To investigate how irregular FISH signals related to cytological fragility at the sites, the frequency of chromosomes showing such signals for each of the BAC probes was quantified (Fig 2A and Supp Fig 2A). This fine-mapping of molecular-level misfolding phenotype (i.e. irregular FISH signals) revealed that the frequency of these signals had a similar distribution along the CFS regions as the cytological breaks. A more extensive analysis at FRA1C and FRA4F indicated that the frequency of misfolding extended beyond the region most affected by cytogenetic lesions and abnormal compaction signals was reduced at BACs that did not frequently overlap with breaks. Furthermore, irregular FISH signals were observed at a similar frequency irrespective whether cytological lesions were present or absent (Supp Fig 2C). This analysis revealed that CFS regions, while highly prone to forming cytogenetic abnormalities, are also characterised by an additional level of instability at a molecular level, indicative of a defect in mitotic chromosome condensation.

As molecular misfolding was observed in chromosomes exposed to replication stress, we wanted to determine if such signals were still present in unperturbed cells, which do not show cytogenetic lesions. Signal phenotypes for two BACs at FRA1C and at FRA4F were examined and surprisingly molecular misfolding was elevated at fragile sites compared to control loci even in the absence of replication stress (Figure 2C). This was particularly pronounced in HCT116 cells, where 60% of chromosomes carried disruptions in FRA4F in the absence of replication stress and argues against these folding defects being caused by rare replication events. Conversely, to determine how replication stress affected mitotic condensation at non-CFS regions,the signal phenotypes for two control regions, located on HSA 1q42.3 and 11q13.2 were examined after aphidicolin treatment. The frequency of atypical signals increased at these non-fragile loci in the presence of replication stress, but remained much lower compared to CFS regions (7 to 27% in RPE1, 11% to 28 % in HCT116) but demonstrate that replication stress, per se, can lead to an increase in the frequency of mitotic condensation defects at typical genomic locations.

**Figure 2:**
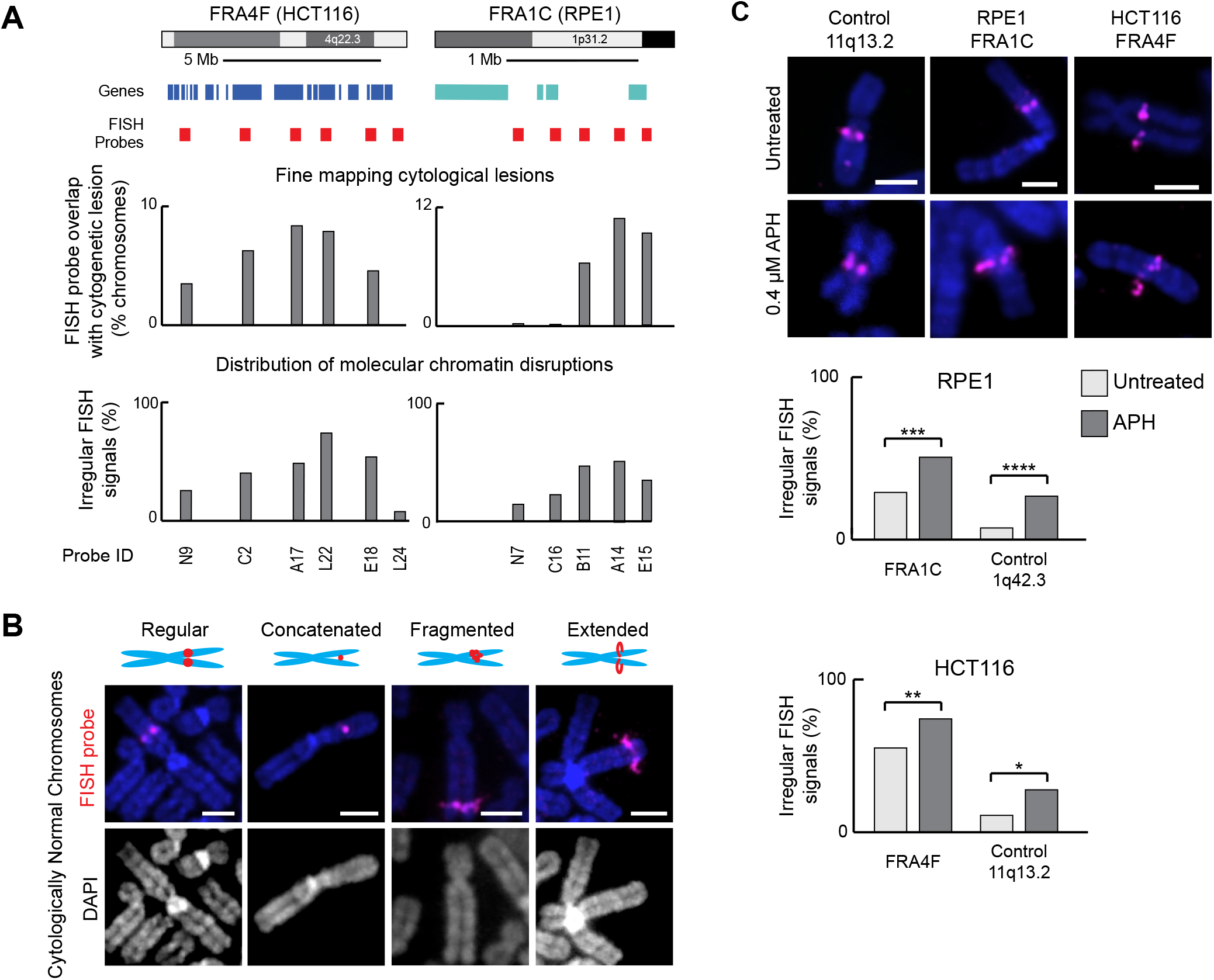
Irregular chromatin structures at CFSs in the presence and absence of replication stress. **A**. Quantification of FISH probe signals across FRA4F (HCT116 cells, n = 439) and FRA1C (RPE1 cells, n = 180) to fine map cytological lesions (top) and distribution of molecular chromatin disruptions (bottom). **B.** Representative irregular FISH probe phenotypes (magenta) at CFS loci on cytogenetically normal chromosomes. Regular - symmetrical, round signals; Concatenated - a single signal sitting between the two sister chromatids; Fragmented - multiple, asymmetric signals; Extended - a signal extending beyond the DAPI stained chromosome area. Scale bar, 10 μm. **C**. Representative chromosomes from untreated cells (top) or cells treated with aphidicolin (APH, bottom), to induce replication stress, hybridised to FISH probes for a non-fragile locus 11q13.2, or fragile loci FRA4F (HCT116 cells) and FRA1C (RPE1 cells). Bottom, quantification of irregular FISH signals in the presence and absence of APH. Scale bar, 10 μm. P-values are for a χ^2^ test.

### Extending G2 reduces cytogenetic lesions and molecular defects at CFS

CFS regions are sites of DNA synthesis on metaphase chromosomes: by utilising a short pulse with the thymidine analogue EdU in mitosis, mitotic DNA synthesis (MIDAS) foci can be observed at cytogenetic CFS lesions (Minocherhomji et al., 2015). To characterise the relationship between MIDAS, cytological lesion formation and molecular misfolding in these cell lines a similar labelling approach was used (Figure 3A). MIDAS occurred at DAPI-faint regions and on many occasions could be seen bridging gaps in chromosomes (Figure 3A and Supplementary Figure 3A). In a subset of metaphases showing extensive damage and wide-spread under condensation, mitotic synthesis foci joined chromosome fragments, overlapping with regions of under-condensation. Consistent with the severity of cytogenetic phenotypes, mitotic synthesis was significantly more frequent in HCT116 than in RPE1 cells: mean number of foci per metaphase was 23.2 and 1.53, respectively (Mann-Whitney U test p < 2.2×10^-16^). Mitotic synthesis was very frequently associated with cytogenetically visible lesions, especially in the HCT116 cell line (91% of EdU foci coincided with lesions, Figure 3A) suggesting that mitotic DNA synthesis preferentially occurs in the context of cytologically under-condensed mitotic chromatin. We also examined the concurrence between molecular-scale misfolding and mitotic synthesis at the FRA4F site by combining FISH with MIDAS labelling. We found that at this site MIDAS rarely appeared on cytogenetically normal regions of the chromosome, even if chromatin at the site showed molecular scale disruptions indicated by an abnormal FISH signal (Figure 3B). Strikingly, this is similar to observations of MIDAS at telomeres, where the fragile telomere phenotype was not found to correlate with the appearance of MIDAS foci (Özer et al., 2018). This observation suggests that unlike cytogenetic disruptions, molecular level misfolding is not accompanied by MIDAS, and raises the possibility that the misfolding phenotypes represent structures that are independent of DNA replication. To assess whether ongoing DNA synthesis was required for chromosome decondensation cells were treated with a high dose of aphidicolin during mitosis. The frequency of cytogenetic lesions did not change, indicating that the mitotic condensation defects are not caused by mitotic synthesis (Supplementary Figure 3B).

Although the structures underlying both cytogenetic and molecular lesions are unclear, we wanted to examine whether they represent intermediates that can be resolved or are permanent defects in mitotic chromatin structure. The duration of G2 following induction of replication stress was artificially prolonged by using the CDK1 inhibitor RO3306, to enable aberrant chromatin structures to be resolved prior to releasing the cells into mitosis (Figure 3C). The frequency of both cytological lesions and molecular misfolding was significantly reduced following RO3306 treatment, indicating that the structures underlying these phenotypes can be subject to replication/repair during G2 (Student’s t-test p < 2.2×10^-16^).

**Figure 3:**
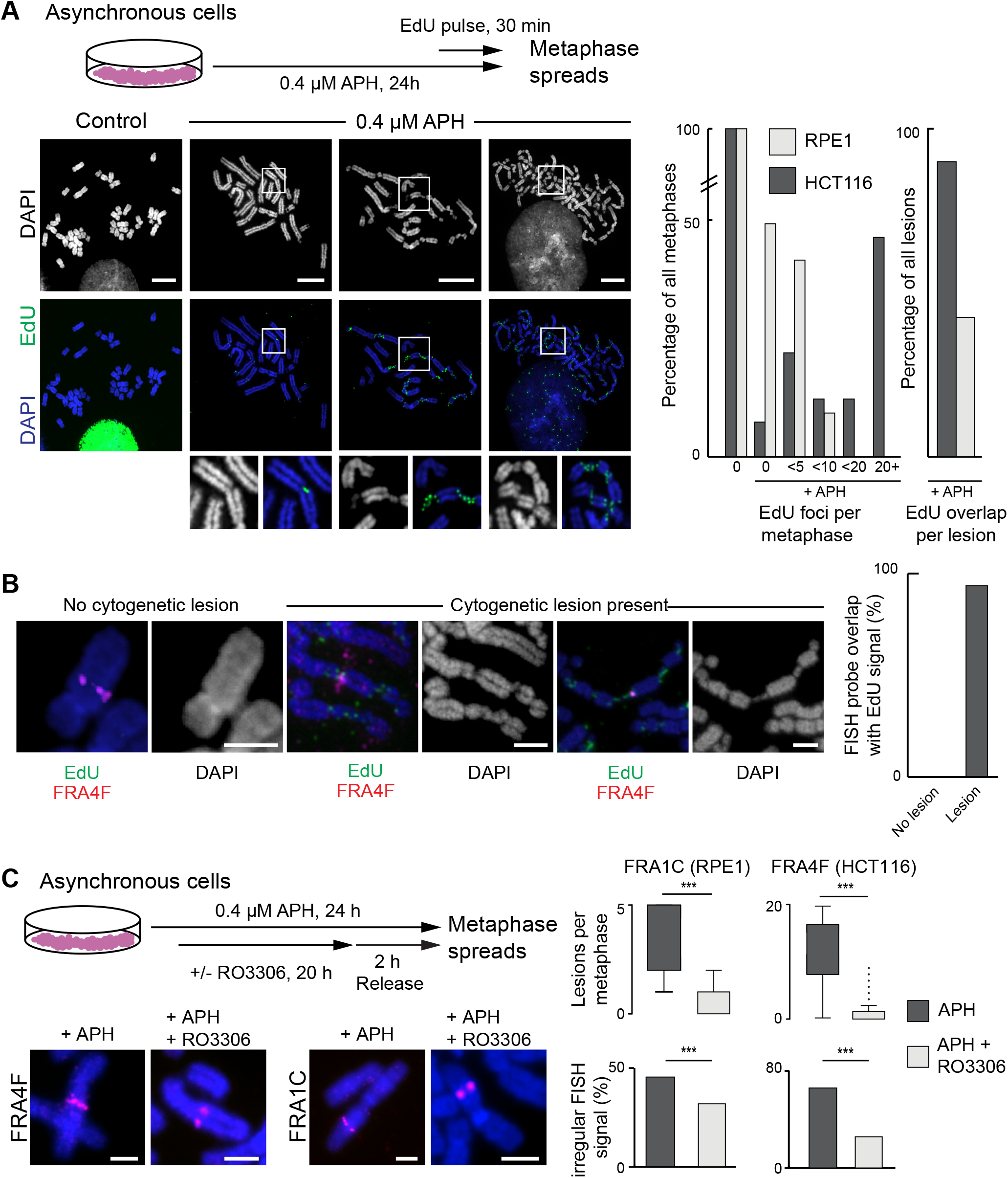
Mitotic DNA synthesis (MIDAS) is coincident with large-scale chromatin disruptions at cytologically visible CFS. **A**. Top, Staining procedure for MIDAS visualised using EdU labelled with FITC-azide. Bottom left, representative metaphase spreads prepared from cells treated with or without aphidicolin. Insets show MIDAS (green signal) and widespread chromosome compaction defects. Right, quantification of MIDAS in RPE1 (n = 65) and HCT116 (n = 82) metaphase spreads from APH treated cells and overlap between cytogenetic lesions and mitotic DNA synthesis in HCT116 (n = 1622 foci) and RPE1 (n = 96 foci) cells. Scale bar, 40 μm. **B**. Atypical FISH signals, cytogenetic lesions and mitotic DNA synthesis (MIDAS) after aphidicolin treatment at the FRA4F locus in HCT116 cells. Representative chromosomes are shown, with increasing degrees of lesions and aberrant condensation. Right, graph showing overlap frequency of FRA4F probe with MIDAS foci on chromosomes in the presence or absence of cytogenetic lesions. Scale bars, 10 μm. **C.** Top, treatment conditions for delaying G2 following induction of replication stress. Bottom left, representative images of FISH signals at the FRA4F (HCT116) or FRA1C (RPE1) loci following aphidicolin treatment followed by a normal duration (left) or extended G2 (right). Right, frequency of cytological breaks (boxplots; P-values are for a Wilcoxon test) and abnormal FISH signals (bar charts; P-values are for a χ^2^ test) after aphidicolin treatment followed by a normal duration (dark grey) or delayed G2 (light grey). Scale bars, 5 μm.

### Chromatin at CFS regions is not remodelled for mitosis

As the mechanism(s) giving rise to aberrant chromatin compaction observed at metaphase (Figure 1A, 2B) were unclear we investigated the possibility that they might arise as interphase chromatin defects. To test this a two-probe FISH approach was used to examine interphase chromatin compaction at two fragile sites: FRA3B (HCT116 cells) and FRA1C (RPE1 cells), in the presence and absence of aphidicolin (Figure 4A) in synchronised cell populations, at different time points throughout the transition from the G1/S boundary through to G2 and mitosis (Figure 4A and Supplementary Figure 4A). No replication-stress induced changes in interphase chromatin structure were observed in FRA3B and FRA1C post-replication, but there was a change in compaction in FRA1C coincident with when the locus replicated in early to mid S-phase. This data indicated that replication-stress *per se* does not induce interphase chromatin structure changes which could explain mitotic condensation failure.

Compaction for mitosis involves many compositional and structural changes, which are required to prepare chromatin for condensation, so we speculated that this process was disrupted at CFS regions. To assess the frequency of misfolding lesions at CFS loci throughout the cell cycle, a premature chromosome condensation assay was used at three CFSs and a control, non-fragile region on HSA 11q13.2. Cells were treated with the phosphatase inhibitor calyculin A, which triggers chromosome condensation, irrespective of cell cycle stage (Figure 4B). This resulted in the formation of prematurely condensed chromosomes with morphologies that are indicative of the cell cycle stage they are derived from: thin and zig-zag shaped in G1; fragmented chromatin in S-phase; cross shaped chromosomes with fuzzy boundaries in G2 cells and typical metaphase chromosomes in mitotic cells (Ono et al., 2013; El Achkar et al., 2005). To assess the condensation capacity of FRA4F, FRA2I, FRA3B and the control location at different phases of the cell cycle, FISH probes mapping to the three locations were hybridised and the morphology of the FISH signals were scored for the different cell cycle stages (G1, S, G2 and M) in the absence of replication stress. To verify the accuracy of our approach, we quantified the frequencies of one-spot (unreplicated) and two-spot (replicated) signals throughout the different cell cycle stages and found that, as expected, one-spot signals were more common at the early stages of the cell cycle and two-spot signals were more frequent in G2 and mitotic chromosomes, especially at the control location (Supplementary Figure 4B). The analysis revealed contrasting dynamics in chromatin competence for condensation at the CFS sites and the control region (Figure 4B). At the control locus, the frequency of atypical signals decreased in the later phases of the cell cycle, with only a small proportion of signals retaining the misfolded phenotype in G2 and M chromosomes. This indicated that chromatin at the locus acquires competency for mitotic compaction as the cell cycle proceeds. In contrast, at the three CFSs, the atypical FISH signals persisted throughout the cell cycle and remained high in mitotic chromosomes, indicating that the process which allows genomic locations to remodel their chromatin environment and compact for mitosis may be disrupted at CFSs. These results also indicated that the molecular lesions manifested at CFSs in mitosis (Figure 2B) are initiated at earlier cell cycle stages.

**Figure 4:**
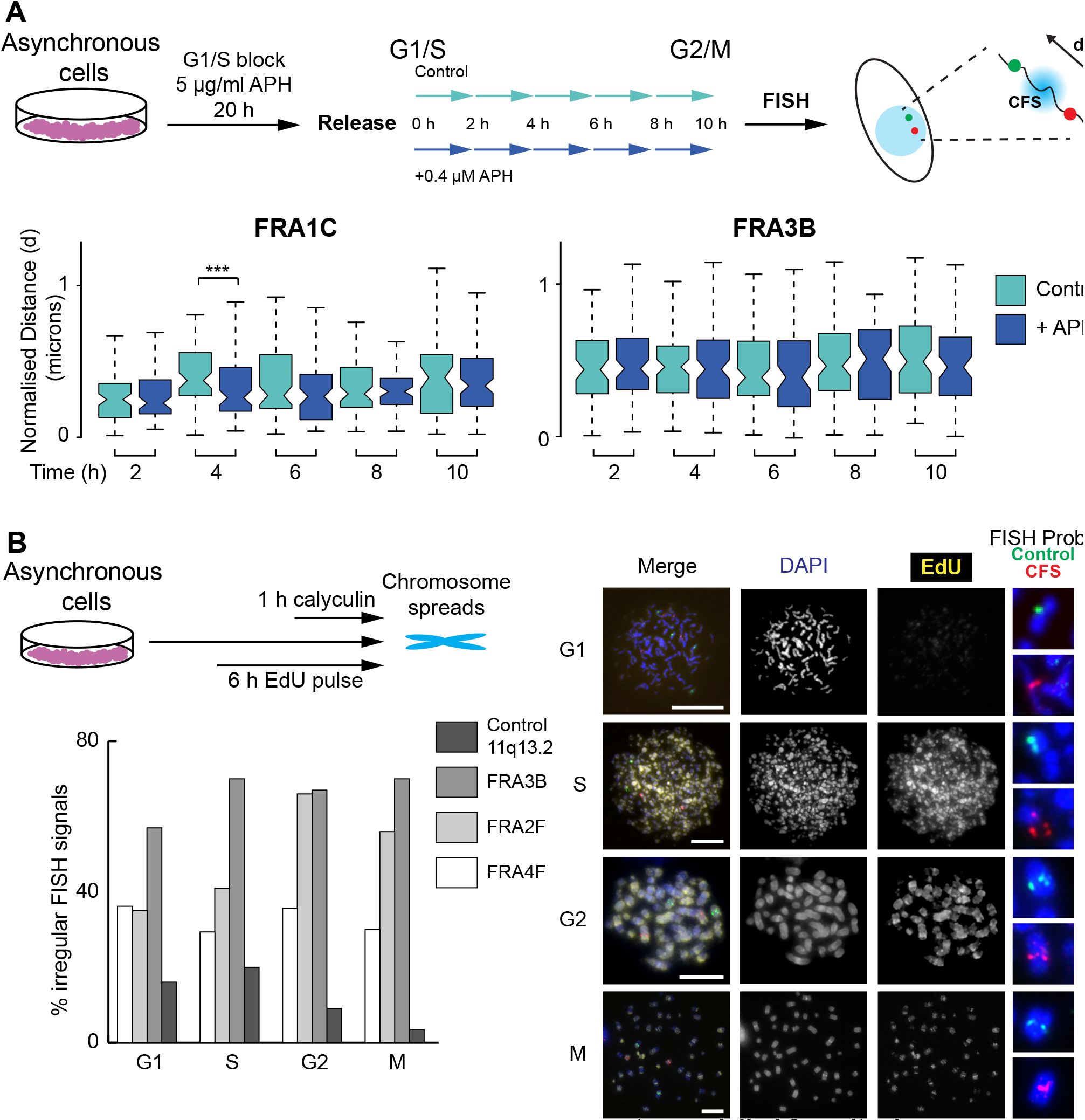
Molecular chromatin disruptions at CFS are not remodelled for mitosis. **A.** Top, model detailing experimental strategy to analyse chromatin folding at CFS regions. Cells progressing synchronously through the cell cycle were harvested every two hours. Bottom, boxplot of normalised inter-fosmid distances between pairs of probes flanking FRA1C (n > 60 RPE1 nuclei) or FRA3B (n > 60 HCT116 nuclei). P-values are for a Wilcoxon test. **B.** Depiction of premature chromosome condensation (PCC) assay (see methods) in HCT116 cells. Cells labelled with EdU (6 h) were condensed using calyculin (1 h), harvested and hybridised to FISH probes for a control locus (11q13.2) and CFSs (FRA3B, FRA4F and FRA2I). Right, representative chromosomes and bottom left, quantification of irregular FISH probe signals at FRA3B (n = 122 chromosomes), FRA2I (n = 136), FRA4F (n = 119) and 11q13.2 control probe (n = 116).

### Cytologically observed defective condensin loading at CFS regions

We considered the potential mechanisms that may affect mitotic compaction at CFS loci. Previous work in yeast has indicated that recruitment of condensin is blocked at slow replication zones to prevent break formation (Hashash et al., 2012) and in mammalian cells condensin recruitment is abrogated after DNA damage (Zhang et al., 2016). To determine if similar processes are applicable here we examined condensin localisation to cytogenetic breaks at CFS loci. Using an antibody against SMC2, a component of both condensin complexes active in mammalian cells, CFS cytogenetic lesions were frequently found to be depleted of condensin (Figure 5A, Supplementary Figure 5A, 5B). Furthermore, the region of condensin depletion appeared to encompass a larger area than the cytogenetic break. On a very small proportion of chromosomes, large regions of SMC2 depletion could be observed in the absence of a cytogenetic break, and at cytogenetic locations that were consistent with frequent CFSs, such as FRA1C (Supplementary Figure 5B). To verify that regions of SMC2 depletion in the absence of cytogenetic abnormalities occur at CFSs, SMC2 immunofluorescence was combined with FISH using probes for the FRA1C and FRA4F CFS regions and confirmed that SMC2 depletions occur at CFS regions (Figure 5B, Supplementary Figure 5C). Consistently, staining for the H3 serine 10 phosphorylation mark, which is acquired on chromatin in preparation for mitotic folding, also showed a depletion at FRA1C on cytogenetically intact mitotic chromosomes (Figure 5C).

As condensin phosphorylation by Cdk1 (Abe et al., 2011) and Chk2 (Zhang et al., 2016) is necessary for chromosome compaction, we speculated that failure of condensin loading at CFSs could be triggered by ATM or ATR signaling, particularly as inhibition of the ATM-Chk2 pathway restores SMC2 association with mitotic chromosomes after Adriamycin induced DNA damage (Zhang et al., 2016). ATM inhibition in the presence of replication stress caused an increase in cytogenetic lesions per metaphase chromosome and an increase in MIDAS (Supplementary Figure 5D). Concomitantly, there were more SMC2 lesions indicative of a role for ATM in registering damage but SMC2 lesions were still present suggesting that the ATM-Chk2 pathway was not responsible for blocking condensin loading after replication stress.

As ATR has a critical role in maintaining replication-dependent genome stability (Cimprich and Cortez, 2008; Zeman and Cimprich, 2014) we also analysed chromosome architecture after replication stress in the presence of ATR inhibitor. Consistent with previous studies (Casper et al., 2002; Durkin et al., 2006) inhibition of the ATR-Chk1 axis caused widespread chromosome shattering (Supplementary Figure 5E). This phenotype resembled calyculin-induced chromosome condensation in replicating nuclei (Figure 4B) so could be due to chromosome condensation prior to complete replication. Immunostaining indicated SMC2 was not correctly recruited to the chromosomes but DNA synthesis was apparent (Supplementary Figure 5E). Nevertheless, condensin depletion appeared to be relevant for repair processes at CFSs; MIDAS labelling with SMC2 immunostaining revealed that DNA synthesis only occurs in regions of mitotic chromosomes depleted of condensin, indicating that uncondensed chromatin maybe a necessary condition for mitotic DNA synthesis (Figure 5D).

**Figure 5:**
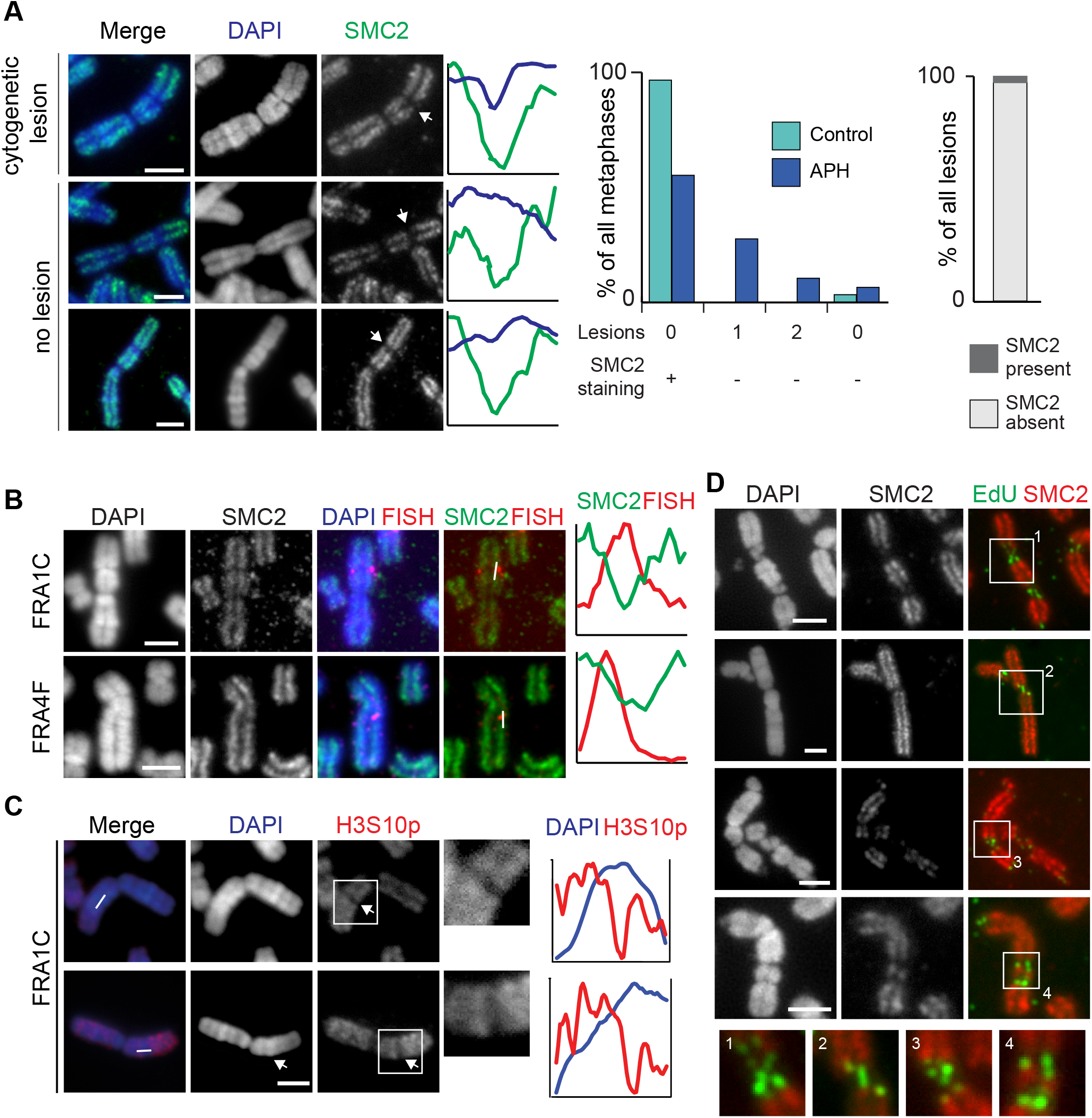
SMC2 depletion at CFSs on mitotic and interphase cells. **A.** Representative images of mitotic chromosomes from RPE1 cells showing regions of SMC2 depletion at cytogenetic lesions (top) and on cytogenetically normal chromosomes (bottom) with intensity profiles of DAPI and SMC2 (regions of interest are marked in Supplementary Figure 5A). Scale bars, 2.5 μm. Right, frequency of cytogenetic lesions and lesion-free SMC2 depletion in the presence or absence of aphidicolin. Far right, quantification of SMC2 occupancy at cytogenetic lesions. **B.** Representative immuno-FISH images showing immunofluorescence staining for SMC2. FISH probes for the FRA1C (RPE1 cells, top) or FRA4F (HCT116 cells, bottom) sites overlap with regions of SMC2 depletion on metaphase chromosomes after replication stress. Right, intensity profiles across the region of interest marked by white line. Scale bars, 2.5 μm. **C.** Representative immunofluorescence staining in RPE1 cells for H3S10 phosphorylation on mitotic chromosomes at the FRA1C locus (arrow) after replication stress. Scale bars, 5 μm. Inset shows enlarged area marked by white box. Right, intensity profiles across the regions of interest indicated by white line. **D.** Representative images of EdU incorporation marking MIDAS on chromosomes from HCT116 cells costained for SMC2. Scale bars, 5μm. Inset shows enlarged area marked by white box. **E.** Top, diagram detailing experimental strategy to analyse SMC2 (condensin I and II) binding by ChIP-seq in HCT116 cells at CFS regions in synchronised cell populations. Bottom, line graphs showing mean normalised condensin binding in 1kb windows at the FRA3B and 4q32.2 sites at different time points in the absence (green) or presence (blue) of aphidicolin.

### Condensin I depletion causes mitotic folding defects at non-fragile locations

As condensin depletion correlates to CFS misfolding in mitosis (Figure 5), we sought to examine the effects of global depletion of the condensin complexes. Initially condensin loss was examined in an HCT116 cell line in which both copies of SMC2 were fused to an AID tag (Supplementary Figure 6A). The HCT116-SMC2-AID cell line showed severe defects in mitotic chromosome structure upon SMC2 degradation: individual chromosomes could not be distinguished and metaphases appeared as a mass of condensed fragments, as described previously (Green et al., 2012). MIDAS foci could still be observed in SMC2 depleted metaphases, indicating condensin is not required for mitotic synthesis (Supplementary Figure 6B), whilst FISH showed a significant increase in molecular misfolding at both control and fragile sites (Supplement Figure 6C).

To explore the different roles of the condensin complexes siRNAs against CAP-H and CAP-D3 were used to deplete condensin I and condensin II complexes, respectively (Supplementary Figure 6D). As for SMC2 depletion (Supplementary Figure 6B) depletion of either condensin complex resulted in mitotic chromosome defects (Green et al., 2012); condensin II depleted chromosomes had a pronounced wavy appearance. Scoring of cytogenetic lesions (Figure 6A) and MIDAS foci (Supplementary Figure 6E) in these chromosomes revealed that although CAP-H and CAP-D3 depletion did not induce lesion formation in unperturbed conditions, there was a significant increase in the frequency of CFS lesions in condensin-depleted chromosomes once replication stress was induced. This is indicative of both replication stress and aberrant condensin loading being necessary for lesion formation at CFS.

**Figure 6:**
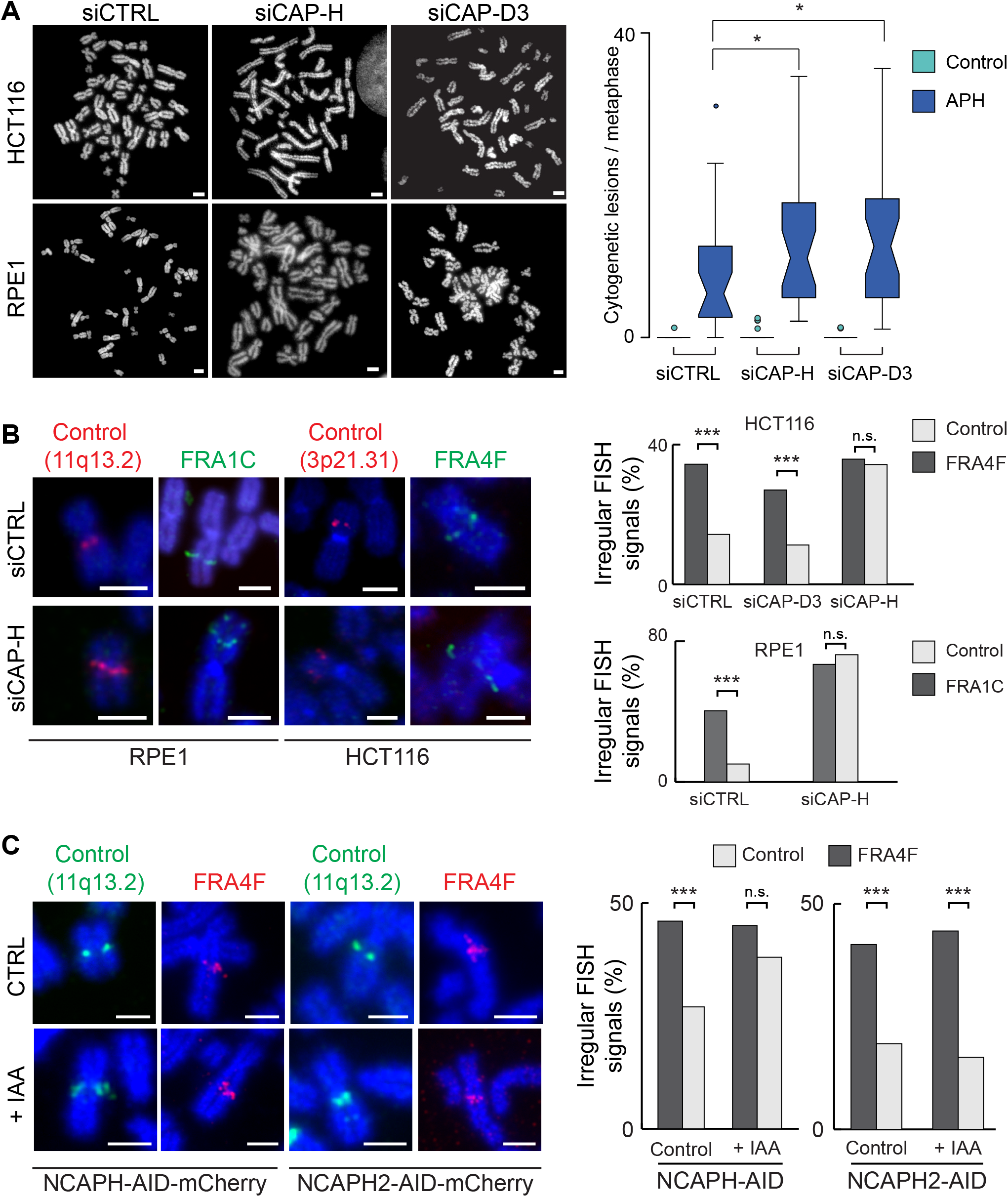
Condensin I depletion causes molecular chromatin lesions in mitosis. **A.** Left, representative images of chromosomal defects in the HCT116 and RPE1 cell lines following depletion of the condensin components CAP-H (condensin I) and CAP-D3 (condensin II). Scale bars, 10 μm. Right, quantification of the number of cytogenetic lesions per metaphase following condensin depletion in the HCT116 cell line in the absence (green) or presence (blue) of aphidicolin. **B.** Left, representative images of chromosomal defects visualised by FISH at fragile (FRA1C and FRA4F) and control loci in RPE1 and HCT116 cell lines following depletion of the condensin components CAP-H and CAP-D3, in the absence of aphidicolin treatment. Scale bars, 5 μm. Right, quantification of abnormal FISH signals, indicative of chromatin disruptions, per metaphase following condensin depletion in the HCT116 and RPE1 cell lines. **C.** Left, representative images showing FISH probes at the fragile FRA4F region and a non-fragile control region (11q13.2) following degradation of the condensin I component CAP-H in HCT116 cells. Scale bars, 5μm. Right, quantification of the frequency of irregular FISH signals, indicative of CAP-H dependent mitotic chromosome misfolding.

We next examined the effect of condensin depletion on the mitotic misfolding phenotype at CFS and control, non-fragile locations by FISH. The frequency of misfolding at CFS locations was significantly higher than control loci in siCTRL-treated cells in both the RPE1 and HCT116 cell line (Figure 6B). However, in cells depleted of CAP-H, the frequency of misfolding was similar across CFSs and control-loci, suggesting that depletion of condensin I is sufficient to recapitulate the misfolding phenotype characteristic of CFS sites at a non-fragile location. In contrast, depletion of CAP-D3 (condensin II) did not affect the frequency of misfolding at non-fragile locations, indicating that the condensin I complex is the primary effector of mitotic misfolding at CFS locations.

To further investigate the role of condensin using an orthogonal approach, we used HCT116 cell lines in which both copies of CAP-H or CAP-H2 were fused to an AID tag, which enabled rapid depletion of either condensin I or II upon addition of auxin (IAA) for 8 hours (Supplementary Figure 6F) (Takagi et al., 2018). Mitotic misfolding, measured by irregular FISH signals, for a control locus at 11q13.2 and the FRA4F fragile site were analysed before and after auxin treatment. As already observed at these sites (Figure 4B) the fragile site locus had a greater extent of mitotic chromosome misfolding in the absence of condensin degradation. However, an increased extent of mitotic chromosome folding was observed for the control locus after CAP-H degradation was triggered (Figure 6C), indicating that defects in condensin I loading underlie mitotic misfolding and that the process is not specific to CFSs but is important genome-wide. In contrast, CAP-H2 degradation did not lead to an increase in misfolding at the control locus.

## Discussion

Replication stress affects genome-wide alterations in individual fork behaviour, leading to activation of extra origins, changes in origin efficiency and potentially, altered replication dynamics (Macheret and Halazonetis, 2018; Courbet et al., 2008). A number of factors can trigger replication stress: oncogene activation, misincorporation of nucleotides or replication-transcription conflicts (Helmrich et al., 2011; Reijns et al., 2015; Hills and Diffley, 2014) but an often overlooked aspect of replication stress is the local chromatin environment. DNA supercoiling (Naughton et al., 2013) or catenanes, paucity of active chromatin marks and unusual DNA structures such as R loops and G-quadruplexes have all been shown to interfere with replication dynamics, suggesting that features of the underlying chromatin environment could be a critical factor linking replication stress to genome instability (Comoglio et al., 2015).

A clear consequence of replication stress is inefficient chromatin compaction and the appearance of cytological lesions at CFSs (Figure 1); we now report a new layer of instability at CFSs, visible at the molecular but not the cytogenetic level (Figure 2). These aberrant structures bear similarity to phenotypes previously seen at telomeres following replication stress and recently, at centromeres (Sfeir et al., 2009). At telomeres, lesions are thought to result from replication problems such as fork collapses or G-quadruplex structures formed by GC-rich telomeric repeats; however, CFSs are not composed of repetitive sequences and it is unclear how small-scale events can lead to fragility and failure of mitotic compaction on such a large genomic scale, suggesting additional factors like chromatin structure or epigenetics play a role as previously proposed (Letessier et al., 2011). The similarity between CFS phenotypes support the idea that mitotic misfolding is a universal feature of CFSs and potentially, other difficult to replicate regions. While classic cytogenetic lesions that characterise CFSs cannot be observed in the absence of aphidicolin, this newly characterised low level of instability, apparent only using molecular techniques such as FISH, is present at these loci at a low frequency even when cells undergo normal replication. This finding indicates the inherent fragility present at CFS regions even in the absence of exogenous replication stress and implicates a model for their instability in physiological contexts such as during tumour development (Alexandrov et al., 2013). Surprisingly our data also shows that after aphidicolin treatment molecular scale lesions are also present at control loci (Figure 2C), indicating a genome-wide link between replication stress and chromatin compaction.

While traditional models envision that CFS instability is mediated primarily through replication dynamics, we suggest that the aberrant processing of post-replicative chromatin, resulting in disrupted mitotic folding, also plays a role. Most genomic regions undergo compositional and structural remodelling, or ‘priming’ of chromatin that facilitates mitotic condensation and sister chromatin separation. This idea is not without precedent: the Kleckner group have suggested that through the cell cycle chromatin is continuously remodelled using the energy of topological stress to drive chromatin compaction (Liang et al., 2015; Kleckner et al., 2004). Mechanistic steps for chromatin folding from interphase to mitosis are poorly defined, but key events include condensin loading, histone H3 phosphorylation and catenane resolution, by topoisomerases. Our data indicates that CFSs are inefficiently primed and remain refractory to compaction: the H3S10p mark and SMC2 are depleted from fragile sites in mitosis (Figure 5) and defective condensin recruitment enhances a range of CFS-specific phenotypes, such as cytogenetic lesions, molecular misfolding and MIDAS (Figure 6 and Supplementary Figure 6).

In addition to inherent low level molecular misfolding (Figure 2), mitotic DNA synthesis (MIDAS) is a frequent feature of CFSs (Figure 3). Chromosome lesions provide a permissive environment for the MIDAS process, and most synthesis occurs in the context of uncondensed chromatin, which is free of condensin (Figure 5D). Our data suggested that SMC2 was not required for MIDAS (Figure 6B) in contrast to a previous study (Minocherhomji et al., 2015). In our study SMC2 was depleted using a degron-based system. This resulted in low levels of SMC2 and a very aberrant chromosome morphology. It has been previously proposed that the MIDAS process constitutes a repair pathway for lesions prior to the completion of mitosis (Minocherhomji et al., 2015; Naim et al., 2013) but our observations suggest that altered chromatin compaction at CFS could also aid their repair. Remarkably, under-condensation in mitosis at unresolved homologous recombination intermediates was found to aid cell division, although these mitotic structures represented a distinct phenotype from CFS lesions (Chan et al., 2018). The primary causes of impaired compaction at CFS are complex: our data suggests that replication or repair intermediates, or aberrant chromatin structures resulting from them, impair condensin recruitment. Consistently extending G2 to allow for the repair of intermediates, results in reduction of both cytogenetic abnormalities and mitotic misfolding (Figure 3C).

Chromosome misfolding is not restricted to CFSs and normal genomic regions show a low frequency of lesion formation in the presence of replication stress suggesting that common fragile sites do not have a unique set of chromatin features (Figure 2). Instead, CFSs are at the extreme end of a spectrum of aberrant chromatin structures that have a propensity to exhibit replication stress and inefficient priming leading to misfolding in mitosis and subsequent chromosome instability if not repaired. Counterintuitively cytological lesions may provide a chromosomal environment to facilitate processes like MIDAS to repair chromatin.

## Acknowledgements

We thank Masatoshi Takagi for condensin degron cell lines and Bernie Ramsahoye, Andrew Wood and Ian Adams for useful comments on the manuscript. We would also like to thank Lizzie Freyer for FACS technical support. This work was supported by the UK MRC (MC_UU_00007/13) and an MRC Studentship to L.B., N.G. is an MRC Senior Research Fellow (MR/J00913X/1).

## Author contributions

L.B., R.N. and C.N, undertook experiments, N.G., L.B., R.N. and C.N. designed experiments and analysed data, K.S. and W.C.E. designed experiments and provided reagents. N.G. and L.B. wrote the manuscript with input from all authors.

## Competing Financial Interests

The authors declare no competing financial interests

## Methods

### Cell culture and transfections

RPE1 and HCT116 cells were cultured in Dulbecco’s Modified Eagle Medium F12 (Gibco, Cat No. 12500-062), supplemented with 10% foetal calf serum, 1% Pen-Strep and 1% L-glutamine. Additionally, growth media for RPE cells also contained 0.3% (w/v) Sodium Bicarbonate (Sigma, Cat. No. S5761). All cells were maintained at 37 °C in an atmosphere of 5% CO2. HCT116 degron cell lines were grown in McCoy’s 5A medium (Gibco, Cat No. 26600-023) supplemented with 10% foetal calf serum and 3 mM L-glutamine. All cell lines were subjected to regular mycoplasma testing. Vectors were transfected into cells using Lipofectamine 2000 (Invitrogen, Cat. No. 11668-019) and Opti-MEM Reduced Serum Medium (Invitrogen, Cat. No. 31985-070). For each transfection 1 μg of construct DNA was mixed with 400 μl Opti-Mem and 5 μl Lipofectamine-2000. To avoid aggregation of DNA and Lipofectamine-2000, the DNA was pre-mixed in 200 μl of Opti-Mem and separately, the 5 μl of Lipofectamine were mixed into 200 μl of Opti-mem. The two components were then mixed together and incubated for 20 min at RT. This transfection mixture was added to the tissue cultures in 2 ml of antibiotic-free media.

### Protein gels and western blotting

Cells were suspended in NuPAGE LDS sample buffer (ThermoFisher) with 10 mM DTT, incubated at 100°C for 5 min and sonicated briefly. Protein samples were resolved on 8% bis-tris gels (ThermoFisher) and transferred to Immobilon-P PVDF 0.45 μm membrane (Merck Millipore) by wet transfer. Membranes were probed with antibodies using standard techniques and detected by enhanced chemiluminescence. Antibodies used for western blotting were as follows: CAP-H (Bethyl A300-603A, 1:1000), CAP-D3 (Bethyl, A300-604A, 1:1000) and GAPDH (Cell Signaling 2118L, 1:5000).

### Cell cycle synchronisation and replication stress induction

Cells were synchronised at the G1/S boundary by addition of high dose aphidicolin (APH, Calbiochem). Media containing 5 μg/ml APH was added to cells for 2 h to block cell cycle and retain cells at the G1/S boundary. Cells were washed in PBS and released in normal growth media. FACS analysis and immunofluorescence of cell populations at 2 h – 10 h following release showed that cells progressed synchronously from S-phase into G2. Replication stress was induced by low dose treatment of APH (0.4 μM APH), for extended periods (12 – 24 h).

### Fluorescence in situ hybridization (FISH)

Probes used in this study are listed below. After mapping fragile sites, the following probes were used to interrogate genomic loci by FISH (probe ID: A14, A17, K5, C2, L12, A13, P21, C14). DNA was prepared from the BACs or Fosmids and labelled as previously described (Naughton et al., 2010). Probes were labelled using a nick translation reaction with the uridine analogues biotin-16-dUTP (Roche, CatNo 11093070910) or digoxigenin-11-dUTP (Roche, CatNo 11093088910). Nick translation was performed in a 20 μl reaction volume, containing 1-1.5 μg DNA with 5 μl each of 0.5 mM dATP, dCTP and dGTP and either 2.5 μl of 1 mM biotin-16-dUTP or 1 μl of 1 mM digoxigenin-11-dUTP. DNase I (Roche, Cat No 4716728001) was added to a final concentration of 1 U/ml and DNA polymerase I (Invitrogen, Cat No 18010025) was added to a final concentration 0.5 U/μl. The reaction was performed in 1 x nick translation salts (NTS) buffer, containing 50 mM Tris pH7.5, 10 mM MgSO4, 0.1 mM DTT and 50 μg/ml BSA for 90 min at 16°C. Unincorporated nucleotides were removed by gel filtration of the NTS reaction through a G50 Sephadex spin column (Roche, Cat No G50DNA-RO). Slides, containing either MAA-fixed chromosome spreads or PFA-fixed nuclei, were treated with 100 μg/ml RNaseA (Invitrogen, Cat No 12091039) in 2 x SSC for 1 h at 37°C, washed briefly in 2 x SSC and dehydrated through an ethanol series (2 min each in 70%, 90% and 100% ethanol). Slides were air dried and baked at 70°C for five min before denaturing. Denaturation was performed in 70% formamide (v/v) in 2x SSC (pH 7.5). Slides containing MAA-fixed chromosome spreads were denatured at 70°C for 1 min, while slides on which cells were cultured and then fixed in 4% PFA were denatured at 80°C for 20 min. Following denaturation, slides were submerged in ice-cold 70% ethanol for 2 min and then dehydrated through 90% and 100% ethanol for 2 min each at RT. For hybridisation, 150 ng of labelled probe was combined with 5 μg of salmon sperm and 10 μg of human Cot1 DNA (Invitrogen, Cat No 15279011). Two volumes of ethanol were added and the probe mix was collected by centrifugation and dried. Dried probes were resuspended in 10 μl of hybridisation buffer containing 50% formamide (v/v), 1% Tween-20 and 10% dextran sulphate (Sigma Aldrich, Cat No D8906-100G) in 2 x SSC. Probes were denatured at 70°C for 5 min and reannealed at 37°C for 15 min and chilled on ice. Probes were pipetted onto slides and hybridisation was performed at 37°C overnight. Coverslips were then removed and slides were washed four times in 2 x SSC at 45°C for 3 min and four times in 0.1 x SSC at 60°C for 3 min. Slides were then blocked in 5% milk in 4 x SSC for 5 min at RT Detection of biotin label was performed with sequential layers of fluorescein (FITC)-conjugated avidin, biotinylated anti-avidin and a further layer of FITC-avidin. Digoxigenin was detected with sequential layers of Rhodamine-conjugated anti-digoxigenin and Texas-Red (TR) –conjugated anti-sheep IgG. Slides were DAPI stained, mounted in Vectashield and imaged on a Zeiss epifluorescence microscope using a 100x objective. Data was collected using micromanager software and analysed using custom scripts in iVision or ImageJ.FISH probes used in this study (all coordinates hg38):

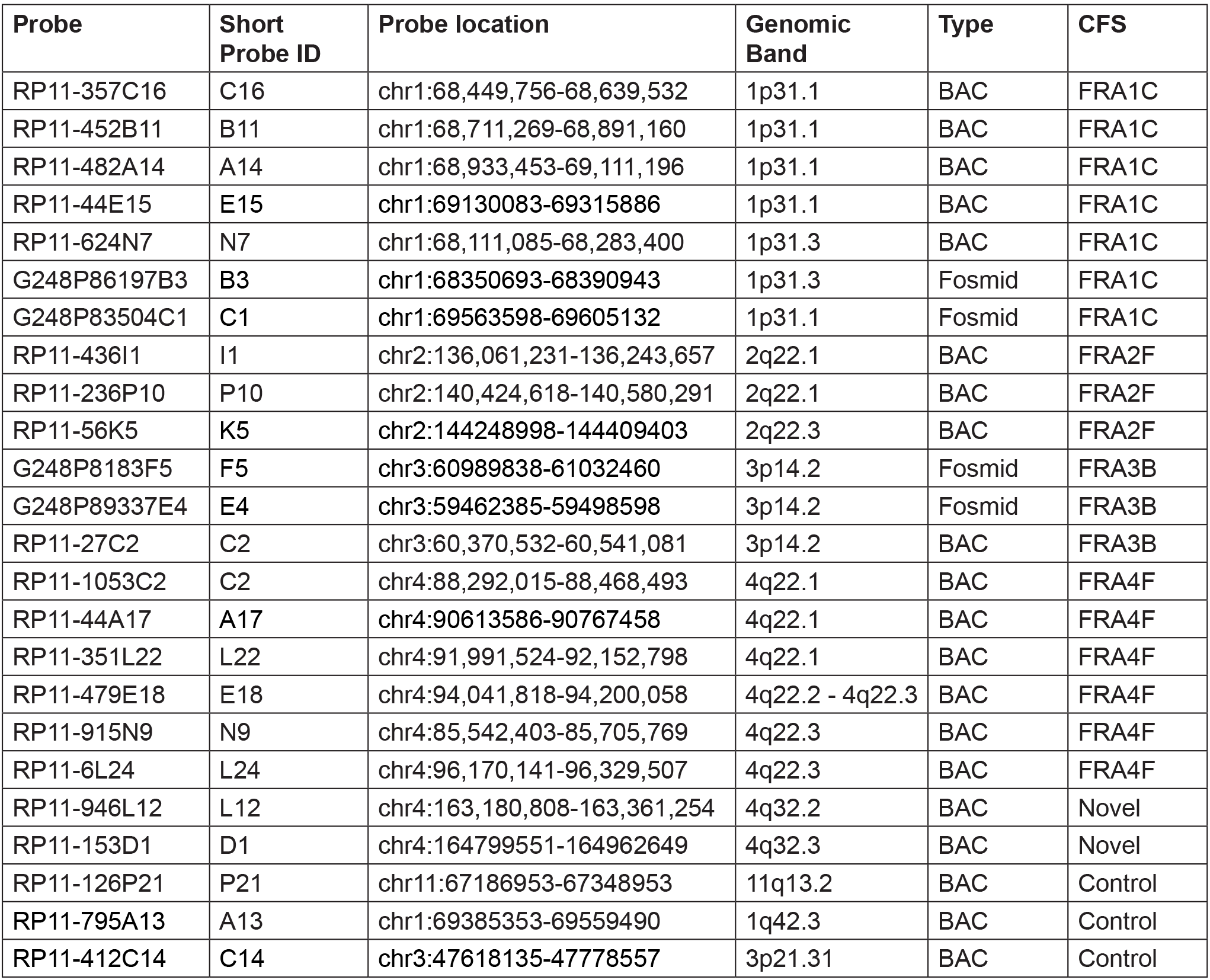

### Preparation of human metaphase chromosomes

RPE1 cells were treated with 0.1 μg/ ml colcemid (Life Technologies, Cat No 15210-040) for 1 h prior to harvest, and HCT116 for 30 min to induce mitotic arrest and increase the number of mitotic cells. Cells were trypsinised and washed in PBS. Hypotonic solution, containing 75 mM KCl was added drop wise to a final 5 ml volume. Hypotonic treatment was performed at RT for 10 min, after which cells were pelleted by centrifugation at 1200 rpm for 5 min and fixed three times in 5 ml of freshly prepared solution of 3:1 ratio (v/v) methanol: acetic acid. The MAA fixative was added to the cell pellet dropwise with constant agitation. Chromosome preparations were stored at −20°C. To prepare slides with metaphase spreads, metaphase chromosome preparations were dropped onto glass slides. The glass slides were pre-treated in a dilute solution of HCl in ethanol for at least an hour prior to use. The chromosome preparations were pelleted by centrifugation at 1500 rpm for 5 min and resuspended in freshly prepared MAA solution until the suspension became cloudy. Two drops of the suspension were dropped onto a pre-treated glass slide from a height of 20 cm and dried at RT overnight before staining or hybridisation.

### Cytogenetic analysis of common fragile sites

To map the location of fragile sites two complementary approaches were used. Firstly, a visual inference of the fragile locus position was made using reverse DAPI banding. Secondly, the position of the fragile site was determined by calculating the distance along the chromosome arm. In this ratio-based approach, the total length (a), in pixels, of the chromosome arm that the break occurred on and the pixel length of the distance between the centromere and the break (b) were measured. The ratio (b) / (a) was calculated and used on scaled models of banded chromosomes (from the International System for Human Cytogenetic Nomenclature) to infer genomic locations for the breaks. The ratios clustered along the chromosome arms, indicating recurrent breaks at CFS locations and the mid-point of each cluster was taken as a putative CFS location. However, as fixation and spreading of chromosomes is likely to cause some distortion, molecular fine-mapping of the most frequent CFS regions was also undertaken using FISH.

### Premature chromosome condensation (PCC) assay

Premature chromosome condensation was induced using the protein phosphatase 1 inhibitor calyculin (Sigma, C5552-10UG). To determine the cell cycle stage of prematurely compacted chromosomes, asynchronously growing HCT116 cells were pulsed with EdU (5 μM) for 6 hours and then treated with 50 ng/ml calyculin for 1 hour. Cells were then harvested using trypsin/versene. As calyculin treatment induced significant cell detachment, media containing the detached cells was collected and centrifuged together with the trypsinised cells. Metaphase spreads were prepared and dropped onto glass slides using the methods described above. Incorporated EdU was labelled in a click reaction by incubating slides with a reaction mixture containing 500 μg/ml CuSO4, 40 μM Alexa Fluor™ 488 Azide (Thermo Fisher Cat. No. A10266) and 20 mg/ml ascorbic acid for 1 hour at RT. Slides were washed three times in PBS for 5 min, stained with DAPI and mounted in Vectashield. Slides were imaged on a Zeiss Epifluorescence microscope using 100x objective. Chromosomal morphology and EdU staining pattern were used to classify chromosomes into G1, S, G2 and M-stage chromosomes.

### Immunofluorescence

For immunofluorescence on metaphase chromosomes, cell suspensions fixed in 3:1 methanol: acetic acid were dropped onto glass slides, allowed to dry incompletely and immediately immersed in PBS for 5 minutes at room temperature. Slides were washed in TEEN buffer (10 mM Triethanolamine-HCl pH 8.5, 2 mM EDTA, 250 mM NaCl) and blocked in 10% fetal calf serum (FCS) at 37°C for 10 minutes. Primary antibodies were added at the required dilutions and incubated in a humidified chamber at 37°C for 30 minutes. Slides were then washed in KB buffer (100 mM Tris-HCl pH 7.7, 1.5 M NaCl, 1% BSA). Secondary antibodies, raised in donkey and conjugated to fluorophores (Jackson Immuno Research), were diluted 1:500 in TEEN buffer, added to the slides and incubated at 37°C for 30 minutes. Slides were washed in KB buffer and stained in 50 μg / ml DAPI for 3 min at RT to detect DNA and nuclei. Slides were mounted in Vectashield (Vector Laboratories, Cat No H-1000) and imaged on a Zeiss Epifluorescence microscope using 100x objective. Primary antibody anti-H3S10p (1:100 dilution, clone CMA313) was a gift from Hiroshi Kimura (Hayashi-Takanaka et al., 2009), and detected with a Texas Red anti-mouse secondary antibody (1:500 dilution, Jackson Immuno Research). Anti-SMC2 antibody was detected with an FITC labebelled anti-rabbit secondary antibody (1:500 dilution, Jackson Immuno Research).

### EdU labelling and detection

EdU was added to exponentially growing cell cultures for 30 min at 5 μM for replication labelling or 20 μM for analysing mitotic DNA synthesis. EdU was detected by incubating slides with a click reaction mixture containing 500 μg/ml CuSO4, 40 μM Alexa Fluor™ 488 Azide (Thermo Fisher Cat. No. A10266) and 20 mg/ ml ascorbic acid for 1 hour at RT. Slides were washed three times in PBS for 5 min, stained with DAPI and mounted in Vectashield. Slides were imaged on a Zeiss Epifluorescence microscope using 100x objective.

### Flow cytometry

For dual EdU and propidium iodide (PI) staining for cell cycle analysis, cells were trypsinised, pelleted and resuspended in PBS at a density of 1.5 x 10^6^ cells/ml. Ethanol was slowly added to the cell suspension to a concentration of 70% to fix and permeabilise the cells, which were incubated on ice for a minimum of 30 min or stored at 4°C for up to 2 weeks. EdU staining was performed using the Click-iT Plus EdU Alexa Fluor 647 Flow Cytometry Assay Kit (Invitrogen, CatNo C10634) following the manufacturer’s instructions. Cells were stained in a solution containing 1 μg/ml PI and 4 μg/ml RNase A in PBS at 2 x 10^6^ cells/ml for a minimum of 30 min at RT Cell cycle analysis was performed on a LSR Fortessa analyser (BD Biosciences) and analysed using FlowJo software.

### HCT116 condensin degron cell lines

SMC2-AID-mClover cells were a derivative of tet-OsTIR1 HCT116 cells established in the Kanemaki laboratory (Natsume et al., 2016). C-terminus targeting constructs for the SMC2 gene contained a 5’ homology arm (410 bp), mAID tag, mClover, resistance cassette and a 3’ homology arm (482 bp) (mAID tag, mClover tag and Hygromycin or G418 resistance cassettes were taken from pMK289 and pMK290. The guide RNA target sequence was TCCACATGTGCTCCTTTGGG. Constructs and resultant cell lines were established using published approaches from the Kanemaki laboratory (Natsume et al., 2016). CAPH-AID-mCherry and CAPH2-AID-mCherry HCT116 cells were a kind gift from the Imamoto lab, RIKEN, Japan (Takagi et al., 2018). For SMC2 degradation SMC2-AID-clover cells were incubated with 1μg/mL doxycycline overnight to induce OsTir1 expression and treated with 500μM Indole-3-acetic acid (IAA) for 24 h. For CAP-H and CAP-H2 degradation, HCT116 cells expressing AID tagged proteins were incubated with 500 μM Indole-3-acetic acid (IAA) for 8 h.

### Computational analysis

All genomic coordinates are HG38. COSMIC mutations were assessed at CFSs by examining the COSMIC track on the UCSC browser.

## Notes

### Competing Interest Statement

The authors have declared no competing interest.

### Summary of Updates

This version of the manuscript clarifies some of the analysis and includes an additional figure panel showing chromosomal fragility after ATM inhibition.

## References

Abe, S., K. Nagasaka, Y. Hirayama, H. Kozuka-Hata, M. Oyama, Y. Aoyagi, C. Obuse, and T. Hirota. 2011. The initial phase of chromosome condensation requires Cdk1-mediated phosphorylation of the CAP-D3 subunit of condensin II. Genes Dev. 25:863–874. doi:10.1101/gad.2016411.

El Achkar, E., M. Gerbault-Seureau, M. Muleris, B. Dutrillaux, and M. Debatisse. 2005. Premature condensation induces breaks at the interface of early and late replicating chromosome bands bearing common fragile sites. Proc. Natl. Acad. Sci. USA. 102:18069–18074. doi:10.1073/pnas.0506497102.

Alexandrov, L.B., S. Nik-Zainal, D.C. Wedge, S.A.J.R. Aparicio, S. Behjati, A. V. Biankin, G.R. Bignell, N. Bolli, A. Borg, A.-L. Børresen-Dale, S. Boyault, B. Burkhardt, A.P. Butler, C. Caldas, H.R. Davies, C. Desmedt, R. Eils, J.E. Eyfjörd, J.A. Foekens, M. Greaves, F. Hosoda, B. Hutter, T. Ilicic, S. Imbeaud, M. Imielinski, M. Imielinsk, N. Jäger, D.T.W. Jones, D. Jones, S. Knappskog, M. Kool, S.R. Lakhani, C. López-Ot\’\in, S. Martin, N.C. Munshi, H. Nakamura, P.A. Northcott, M. Pajic, E. Papaemmanuil, A. Paradiso, J. V. Pearson, X.S. Puente, K. Raine, M. Ramakrishna, A.L. Richardson, J. Richter, P. Rosenstiel, M. Schlesner, T.N. Schumacher, P.N. Span, J.W. Teague, Y. Totoki, A.N.J. Tutt, R. Valdés-Mas, M.M. van Buuren, L. van t Veer, A. Vincent-Salomon, N. Waddell, L.R. Yates, Australian Pancreatic Cancer Genome Initiative, ICGC Breast Cancer Consortium, ICGC MMML-Seq Consortium, ICGC PedBrain, J. Zucman-Rossi, P.A. Futreal, U. McDermott, P. Lichter, M. Meyerson, S.M. Grimmond, R. Siebert, E. Campo, T. Shibata, S.M. Pfister, P.J. Campbell, and M.R. Stratton. 2013. Signatures of mutational processes in human cancer. Nature. 500:415–421. doi:10.1038/nature12477.

Antonin, W., and H. Neumann. 2016. Chromosome condensation and decondensation during mitosis. Curr. Opin. Cell Biol. 40:15–22. doi:10.1016/j.ceb.2016.01.013.

Arlt, M.F., J.G. Mulle, V.M. Schaibley, R.L. Ragland, S.G. Durkin, S.T. Warren, and T.W. Glover. 2009. Replication Stress Induces Genome-wide Copy Number Changes in Human Cells that Resemble Polymorphic and Pathogenic Variants. Am. J. Hum. Genet. 84:339–350. doi:10.1016/j.ajhg.2009.01.024.

Bignell, G.R., C.D. Greenman, H. Davies, A.P. Butler, S. Edkins, J.M. Andrews, G. Buck, L. Chen, D. Beare, C. Latimer, S. Widaa, J. Hinton, C. Fahey, B. Fu, S. Swamy, G.L. Dalgliesh, B.T. Teh, P. Deloukas, F. Yang, P.J. Campbell, P.A. Futreal, and M.R. Stratton. 2010. Signatures of mutation and selection in the cancer genome. Nature. 463:893–898. doi:10.1038/nature08768.

Blin, M., B. Le Tallec, V. Nähse, M. Schmidt, C. Brossas, G.A. Millot, M.N. Prioleau, and M. Debatisse. 2019. Transcription-dependent regulation of replication dynamics modulates genome stability. Nat. Struct. Mol. Biol. 26:58–66. doi:10.1038/s41594-018-0170-1.

Casper, A.M., P. Nghiem, M.F. Arlt, and T.W. Glover. 2002. ATR regulates fragile site stability. Cell. 111:779–789. doi:10.1016/S0092-8674(02)01113-3.

Cha, R.S., and N. Kleckner. 2002. ATR homolog Mec1 promotes fork progression, thus averting breaks in replication slow zones. Science (80-.). 297:602–606. doi:10.1126/science.1071398.

Chan, Y.W., K. Fugger, and S.C. West. 2018. Unresolved recombination intermediates lead to ultra-fine anaphase bridges, chromosome breaks and aberrations. Nat. Cell Biol. 20:92–103. doi:10.1038/s41556-017-0011-1.

Cimprich, K.A., and D. Cortez. 2008. ATR: An essential regulator of genome integrity. Nat. Rev. Mol. Cell Biol. doi:10.1038/nrm2450.

Comoglio, F., T. Schlumpf, V. Schmid, R. Rohs, C. Beisel, and R. Paro. 2015. High-Resolution Profiling of Drosophila Replication Start Sites Reveals a DNA Shape and Chromatin Signature of Metazoan Origins. Cell Rep. 11:821–834. doi:10.1016/j.celrep.2015.03.070.

Courbet, S., S. Gay, N. Arnoult, G. Wronka, M. Anglana, O. Brison, and M. Debatisse. 2008. Replication fork movement sets chromatin loop size and origin choice in mammalian cells. Nature. 455:557–560. doi:10.1038/nature07233.

Durkin, S.G., M.F. Arlt, N.G. Howlett, and T.W. Glover. 2006. Depletion of CHK1, but not CHK2, induces chromosomal instability and breaks at common fragile sites. Oncogene. 25:4381–4388. doi:10.1038/sj.onc.1209466.

Durkin, S.G., and T.W. Glover. 2007. Chromosome fragile sites. Annu.Rev.Genet. 41:169–92.:169–192.

Fungtammasan, A., E. Walsh, F. Chiaromonte, K.A. Eckert, and K.D. Makova. 2012. A genome-wide analysis of common fragile sites: What features determine chromosomal instability in the human genome? 993–1005. doi:10.1101/gr.134395.111.A.

Gibcus, J.H., K. Samejima, A. Goloborodko, I. Samejima, N. Naumova, J. Nuebler, M.T. Kanemaki, L. Xie, J.R. Paulson, W.C. Earnshaw, L.A. Mirny, and J. Dekker. 2018. A pathway for mitotic chromosome formation. Science (80-.). 359. doi:10.1126/science.aao6135.

Gordon, D.J., B. Resio, and D. Pellman. 2012. Causes and consequences of aneuploidy in cancer. Nat. Publ. Gr. 13:189–203. doi:10.1038/nrg3123.

Green, L.C., P. Kalitsis, T.M. Chang, M. Cipetic, J.H. Kim, O. Marshall, L. Turnbull, C.B. Whitchurch, P. Vagnarelli, K. Samejima, W.C. Earnshaw, K.H.A. Choo, and D.F. Hudson. 2012. Contrasting roles of condensin I and condensin II in mitotic chromosome formation. J. Cell Sci. 125:1591–1604. doi:10.1242/jcs.097790.

Hashash, N., A.L. Johnson, and R.S. Cha. 2012. Topoisomerase II-and condensin-dependent breakage of MEC1ATR-sensitive fragile sites occurs independently of spindle tension, anaphase, or cytokinesis. PLoS Genet. 8:e1002978. doi:10.1371/journal.pgen.1002978.

Helmrich, A., M. Ballarino, and L. Tora. 2011. Collisions between Replication and Transcription Complexes Cause Common Fragile Site Instability at the Longest Human Genes. Mol. Cell. 44:966–977. doi:10.1016/j.molcel.2011.10.013.

Hills, S.A., and J.F.X. Diffley. 2014. DNA replication and oncogene-induced replicative stress. Curr. Biol. 24:R435–44. doi:10.1016/j.cub.2014.04.012.

Hosseini, S.A., S. Horton, J.C. Saldivar, S. Miuma, M.R. Stampfer, and N.A. Heerema. 2013. Common Chromosome Fragile Sites in Human and Murine Epithelial Cells and FHIT / FRA3B Loss-Induced Global Genome Instability. 1029:1017–1029. doi:10.1002/gcc.

Kleckner, N., D. Zickler, G.H. Jones, J. Dekker, R. Padmore, J. Henle, and J. Hutchinson. 2004. A mechanical basis for chromosome function. Proc. Natl. Acad. Sci. USA. 101:12592–12597. doi:10.1073/pnas.0402724101.

Letessier, A., G.A. Millot, S. Koundrioukoff, A.-M. Lachages, N. Vogt, R.S. Hansen, B. Malfoy, O. Brison, and M. Debatisse. 2011. Cell-type-specific replication initiation programs set fragility of the FRA3B fragile site. Nature. 470:120–123. doi:10.1038/nature09745.

Liang, Z., D. Zickler, M. Prentiss, F.S. Chang, G. Witz, K. Maeshima, and N. Kleckner. 2015. Chromosomes Progress to Metaphase in Multiple Discrete Steps via Global Compaction/Expansion Cycles. Cell. 161:1124–1137. doi:10.1016/j.cell.2015.04.030.

Lipp, J.J., T. Hirota, I. Poser, and J.-M. Peters. 2007. Aurora B controls the association of condensin I but not condensin II with mitotic chromosomes. J. Cell Sci. 120:1245–1255. doi:10.1242/jcs.03425.

Macheret, M., and T.D. Halazonetis. 2018. Intragenic origins due to short G1 phases underlie oncogene-induced DNA replication stress. Nature. 555:112–116. doi:10.1038/nature25507.

Madireddy, A., S.T. Kosiyatrakul, R.A. Boisvert, E. Herrera-Moyano, M.L. García-Rubio, J. Gerhardt, E.A. Vuono, N. Owen, Z. Yan, S. Olson, A. Aguilera, N.G. Howlett, and C.L. Schildkraut. 2016. FANCD2 Facilitates Replication through Common Fragile Sites. Mol. Cell. 64:388–404. doi:10.1016/j.molcel.2016.09.017.

Minocherhomji, S., S. Ying, V.A. Bjerregaard, S. Bursomanno, A. Aleliunaite, W. Wu, H.W. Mankouri, H. Shen, Y. Liu, and I.D. Hickson. 2015. Replication stress activates DNA repair synthesis in mitosis. Nature. 528:286–290. doi:10.1038/nature16139.

Naim, V., T. Wilhelm, M. Debatisse, and F. Rosselli. 2013. ERCC1 and MUS81-EME1 promote sister chromatid separation by processing late replication intermediates at common fragile sites during mitosis. Nat. Cell Biol. doi:10.1038/ncb2793.

Natsume, T., T. Kiyomitsu, Y. Saga, and M.T. Kanemaki. 2016. Rapid Protein Depletion in Human Cells by Auxin-Inducible Degron Tagging with Short Homology Donors. Cell Rep. 15:210–218. doi:10.1016/j.celrep.2016.03.001.

Naughton, C., N. Avlonitis, S. Corless, J.G. Prendergast, I.K. Mati, P.P. Eijk, S.L. Cockroft, M. Bradley, B. Ylstra, and N. Gilbert. 2013. Transcription forms and remodels supercoiling domains unfolding large-scale chromatin structures. Nat. Struct. Mol. Biol. 20:387–395. doi:10.1038/nsmb.2509.

Negrini, S., V.G. Gorgoulis, and T.D. Halazonetis. 2010. Genomic instability an evolving hallmark of cancer. Nat. Rev. Mol. Cell Biol. doi:10.1038/nrm2858.

Ono, T., D. Yamashita, and T. Hirano. 2013. Condensin II initiates sister chromatid resolution during S phase. 200:429–441. doi:10.1083/jcb.201208008.

Özer, Ö., R. Bhowmick, Y. Liu, and I.D. Hickson. 2018. Human cancer cells utilize mitotic DNA synthesis to resist replication stress at telomeres regardless of their telomere maintenance mechanism. Oncotarget. 9:15836–15846. doi:10.18632/oncotarget.24745.

Reijns, M.A.M., H. Kemp, J. Ding, S.M. de Procé, A.P. Jackson, and M.S. Taylor. 2015. Lagging-strand repli-cation shapes the mutational landscape of the genome. Nature. 1–17. doi:10.1038/nature14183.

Samejima, K., I. Samejima, P. Vagnarelli, H. Ogawa, G. Vargiu, D.A. Kelly, F. de Lima Alves, A. Kerr, L.C. Green, D.F. Hudson, S. Ohta, C.A. Cooke, C.J. Farr, J. Rappsilber, and W.C. Earnshaw. 2012. Mitotic chromosomes are compacted laterally by KIF4 and condensin and axially by topoisomerase II$α$. J. Cell Biol. 199:755–770. doi:10.1083/jcb.201202155.

Sfeir, A., S.T. Kosiyatrakul, D. Hockemeyer, S.L. MacRae, J. Karlseder, C.L. Schildkraut, and T. de Lange. 2009. Mammalian telomeres resemble fragile sites and require TRF1 for efficient replication. Cell. 138:90–103. doi:10.1016/j.cell.2009.06.021.

Sonneville, R., R. Bhowmick, S. Hoffmann, N. Mailand, I.D. Hickson, and K. Labib. 2019. TRAIP drives replisome disassembly and mitotic DNA repair synthesis at sites of incomplete DNA replication. Elife. 8:1–19. doi:10.7554/eLife.48686.

Takagi, M., T. Ono, T. Natsume, C. Sakamoto, M. Nakao, N. Saitoh, M.T. Kanemaki, T. Hirano, and N. Ima-moto. 2018. Ki-67 and condensins support the integrity of mitotic chromosomes through distinct mechanisms. J. Cell Sci. 131:jcs212092. doi:10.1242/jcs.212092.

Le Tallec, B., B. Dutrillaux, A.-M. Lachages, G.A. Millot, O. Brison, and M. Debatisse. 2011. Molecular profiling of common fragile sites in human fibroblasts. Nat. Struct. Mol. Biol. 18:1421–1423. doi:10.1038/nsmb.2155.

Le Tallec, B., S. Koundrioukoff, T. Wilhelm, A. Letessier, O. Brison, and M. Debatisse. 2014. Updating the mechanisms of common fragile site instability: how to reconcile the different views? Cell Mol.Life Sci. 71:4489–4494. doi:10.1007/s00018-014-1720-2.

Le Tallec, B., G.A. Millot, M.E. Blin, O. Brison, B. Dutrillaux, and M. Debatisse. 2013. Common fragile site profiling in epithelial and erythroid cells reveals that most recurrent cancer deletions lie in fragile sites hosting large genes. Cell Rep. 4:420–428. doi:10.1016/j.celrep.2013.07.003.

Wechsler, T., S. Newman, and S.C. West. 2011. Aberrant chromosome morphology in human cells defective for Holliday junction resolution. Nature. 471:642–646. doi:10.1038/nature09790.

Wei, P.C., A.N. Chang, J. Kao, Z. Du, R.M. Meyers, F.W. Alt, and B. Schwer. 2016. Long Neural Genes Harbor Recurrent DNA Break Clusters in Neural Stem/Progenitor Cells. Cell. 164:644–655. doi:10.1016/j.cell.2015.12.039.

Wilson, T.E., M.F. Arlt, S.H. Park, S. Rajendran, M. Paulsen, M. Ljungman, and T.W. Glover. 2015. Large transcription units unify copy number variants and common fragile sites arising under replication stress. 189–200. doi:10.1101/gr.177121.114.Freely.

Zeman, M.K., and K.A. Cimprich. 2014. Causes and consequences of replication stress. Nat. Cell Biol. 16:2–9. doi:10.1038/ncb2897.

Zhang, T., S.L. Si-Hoe, D.F. Hudson, and U. Surana. 2016. Condensin recruitment to chromatin is inhibited by Chk2 kinase in response to DNA damage. Cell Cycle. 15:3454–3470. doi:10.1080/15384101.2016.1249075.

Zhu, J., H.-J. Tsai, M.R. Gordon, and R. Li. 2018. Cellular Stress Associated with Aneuploidy. Dev. Cell. 44:420–431. doi:10.1016/j.devcel.2018.02.002.

