## Supplementary data for "Common fragile sites are characterised by faulty condensin loading after replication stress"

**Supplementary Table 1. Common fragile sites analysed in HCT116 and RPE1 cell lines using DAPI banding and fine-mapping using FISH probes (co-ordinates hg38)**

| CFS Name | Cell type specificity (% of all breaks observed in cell type) | | Genomic Location | Fine-mapping |
|---|---|---|---|---|
| | RPE1 | HCT116 | | |
| FRA1C | 18.6 % | 5.8 % | 1p31.2 | • Fine mapping with BAC probes. Fragility found at a 0.6 Mb region around chr1: 68.7-69.3 Mb |
| Novel | 11.9 % | Not fragile | 4q32.2 | • Fine-mapping with BAC probes. Fragility found to span a 1Mb region overlapping with the MARCH1 gene at 4q32.2-4q32.3 boundary |
| FRA3B | Not fragile | 23.4 % | 3p14.2 | • Fine-mapping with fosmid probes. Fragility localised to a 1 Mb region overlapping with the FHIT gene at 3p14.2 |
| FRA4F | Not fragile | 11.0 % | 4q22.2 | • Fine-mapping with BAC probes. Fragility localised to a 5 Mb region between chr4: 88.3-94.2 Mb |
| FRA2F | 8.5 % | 5.8 % | 2q22.2 | • Fine-mapping with BAC probes. Fragility localised to a 2 Mb region between chr2: 141.4-143.3 Mb |
| FRA3O | 16.9 % | 1.3% | 3q26.31 | |
| FRA7E | 10.2% | Not fragile | 7q21.11 | |
| FRA2I | Not fragile | 17.5 % | 2q33.2 | |
| FRA2T | Not fragile | 9.7 % | 2q24.2 | |
| FRA4C | 6.8 % | Not Fragile | 4q31.1 | |
| FRA7K | Not Fragile | 7.1 % | 7q31.1 | |
| Control | Not fragile | Not fragile | 11q13.2 | |

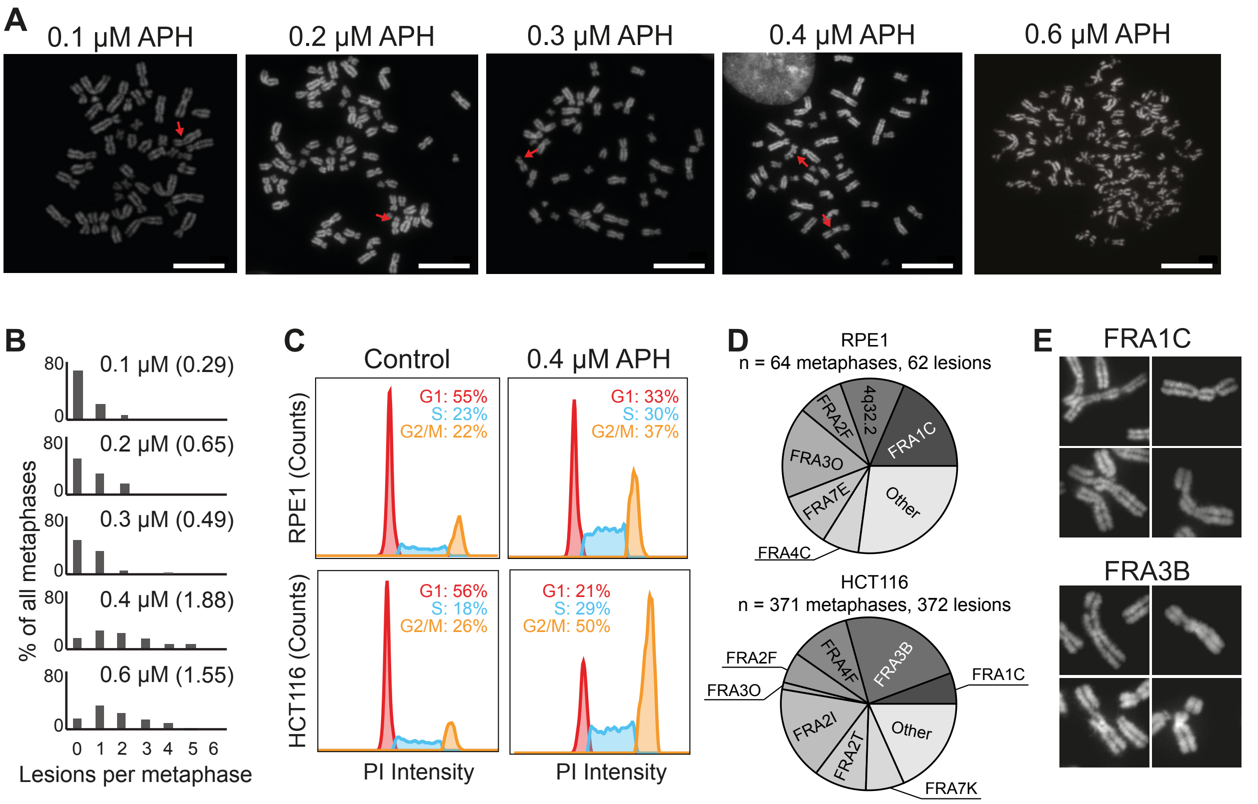


**Supplementary Figure 1: Aphidicolin induced fragility in HCT116 and RPE1 cell lines**

**A**. Representative metaphases following treatment of HCT116 cells with different concentrations of APH for 24 hours. Lesions are indicated by red arrows. Right, metaphase showing multiple regions of incomplete axial compaction following treatment with 0.6 μM APH.

**B**. Proportions of HCT116 metaphase spreads showing different numbers of lesions following treatment with 0.1, 0.2, 0.3, 0.4 or 0.6 μM aphidicolin for 24 hours. A minimum of 52 metaphases were characterised per condition.

**C.** Cell cycle profiles in the HCT116 and RPE1 cell lines in control conditions and following treatment with 0.4 μM APH for 24 hours, analysed via flow cytometry. Proportions of cells in the different stages of the cell cycle.

**D.** Pie charts showing proportion of lesions occurring at specific CFS locations.

**E.** Representative images of lesion morphologies at two different CFSs, FRA1C (top) and FRA3B (bottom).

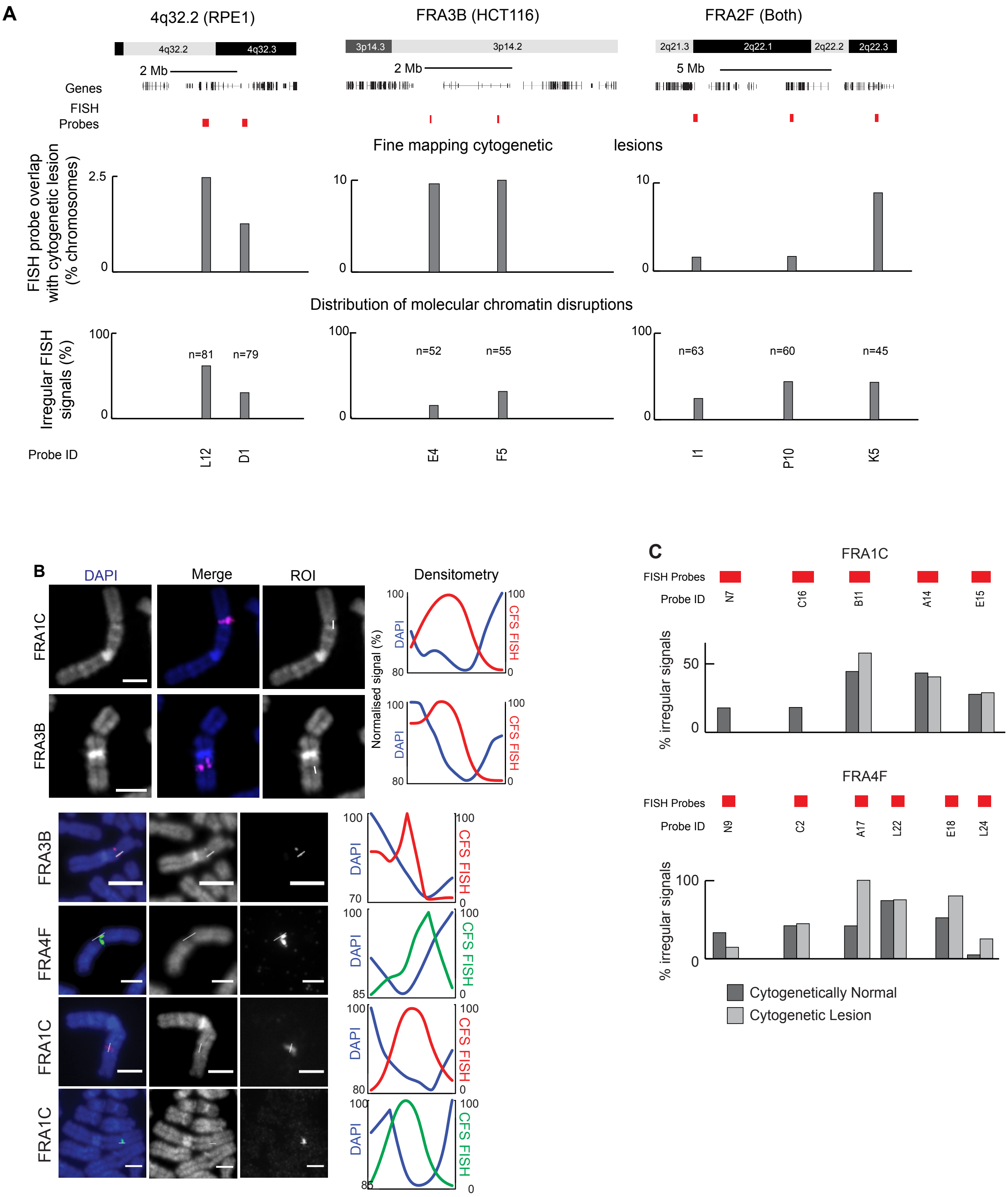


Supplementary Figure 2

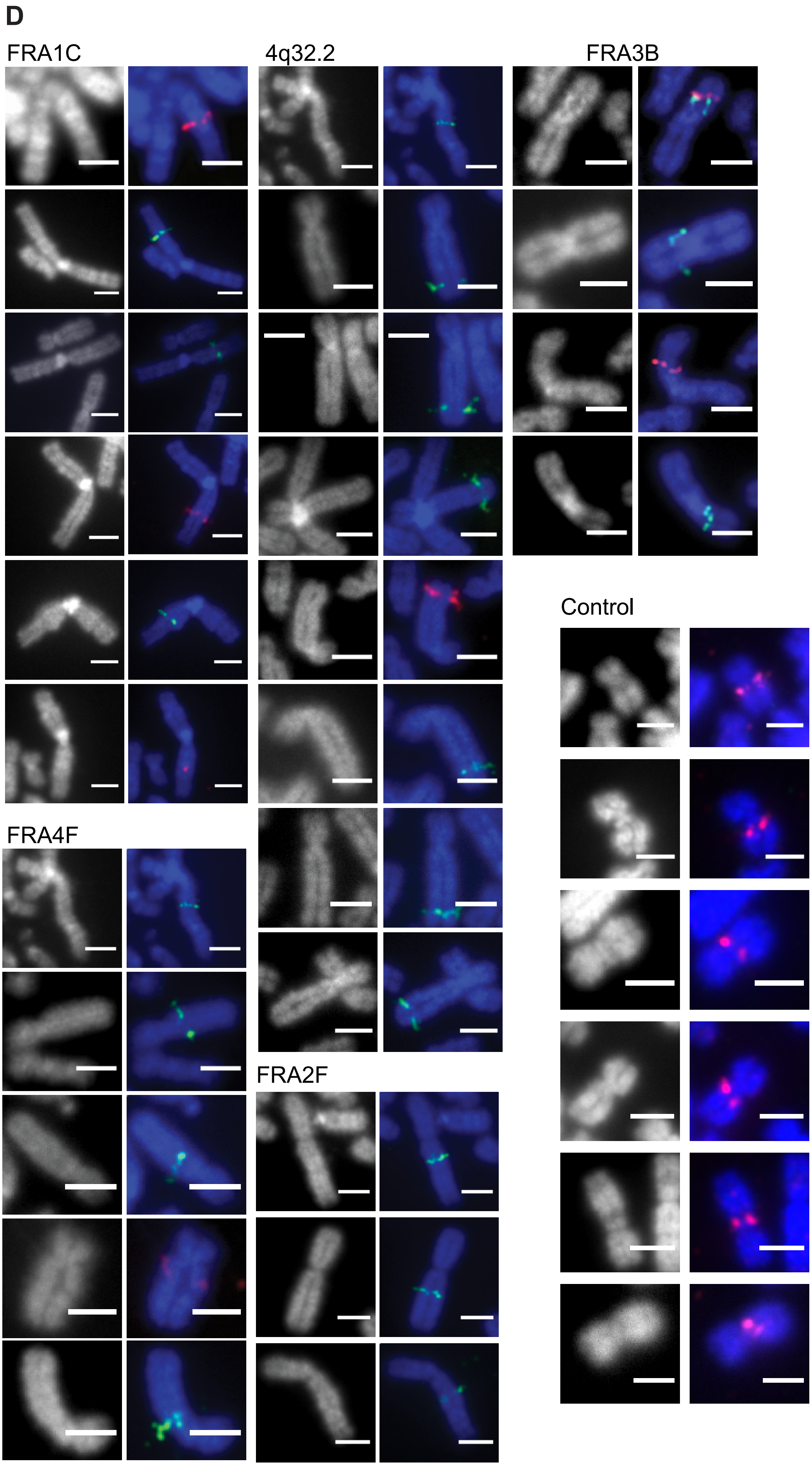

Supplementary Figure 2

**Supplementary Figure 2: CFS FISH probe signals across cytogenetic lesions and cytogenetically normal CFS regions.**

**A**. Top, diagram showing FISH probes (red) spanning lesions at 4q32.2 (RPE1 cells), FRA3B (HCT116 cells) and FRA2F (Both cell lines). Bottom, probes were hybridised to metaphase spreads from cells treated with aphidicolin and counterstained with DAPI followed by quantification to fine map cytological lesions and distribution of molecular chromatin disruptions.

**B.** Representative images showing FISH signals at CFSs FRA1C, FRA2F, 4q32.2, FRA4F and FRA3B. Quantification of FISH signal and DAPI signal marked by white line.

**C.** Quantification of irregular FISH signals at the FRA1C (RPE1) and FRA4F (HCT116) CFS sites in the absence (top) or presence (bottom) of cytogenetic lesions.

**D.** Images showing irregular FISH signals at common fragile sites (FRA1C, 4q32.2, FRA4F, FRA3B) but not control loci.

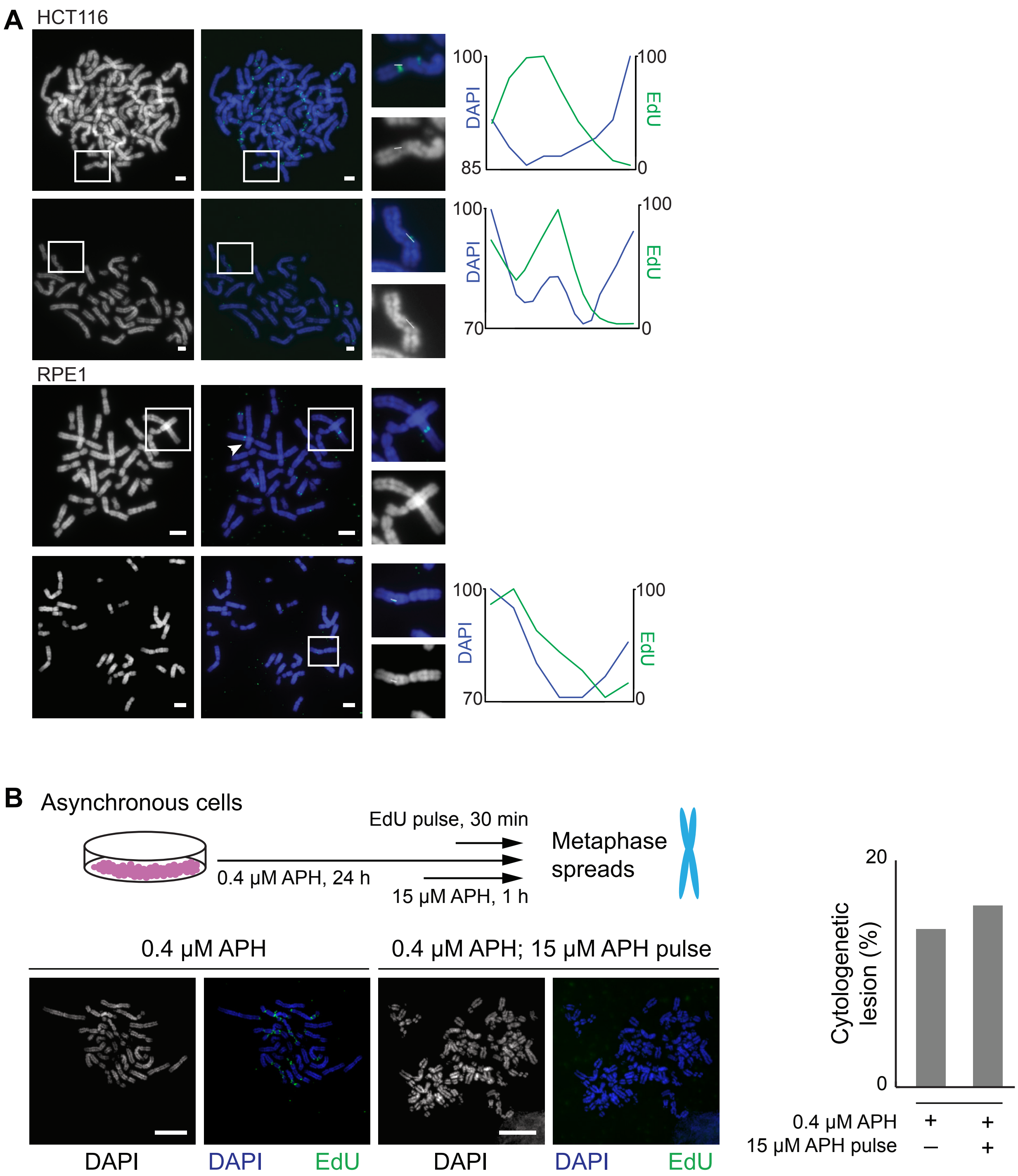


**Supplementary Figure 3: MIDAS and chromatin decondensation following replication stress.** **A**. Representative images showing mitotic DNA synthesis in HCT116 and RPE1 cell lines following 24 h treatment with 0.4 μM APH. Inset, selected cytogenetic lesions with mitotic DNA synthesis. Right, intensity profiles of mitotic synthesis foci across the cytogenetic lesions. White lines indicate the regions selected for the intensity profile, produced in ImageJ.
**B.** Schematic to assess relationship between MIDAS and aberrant chromosome compaction. Cells were exposed to low concentration aphidicolin to induce replication stress and then high dose aphidicolin to inhibit DNA synthesis. Bottom left, representative metaphases showing MIDAS (EdU, green) in cells treated only with low dose aphidicolin and those treated with additional high dose aphidicolin to inhibit MIDAS. Right, quantification of metaphases showing chromosomal lesions in the presence or absence of transcription. Scale bar, 40μm.

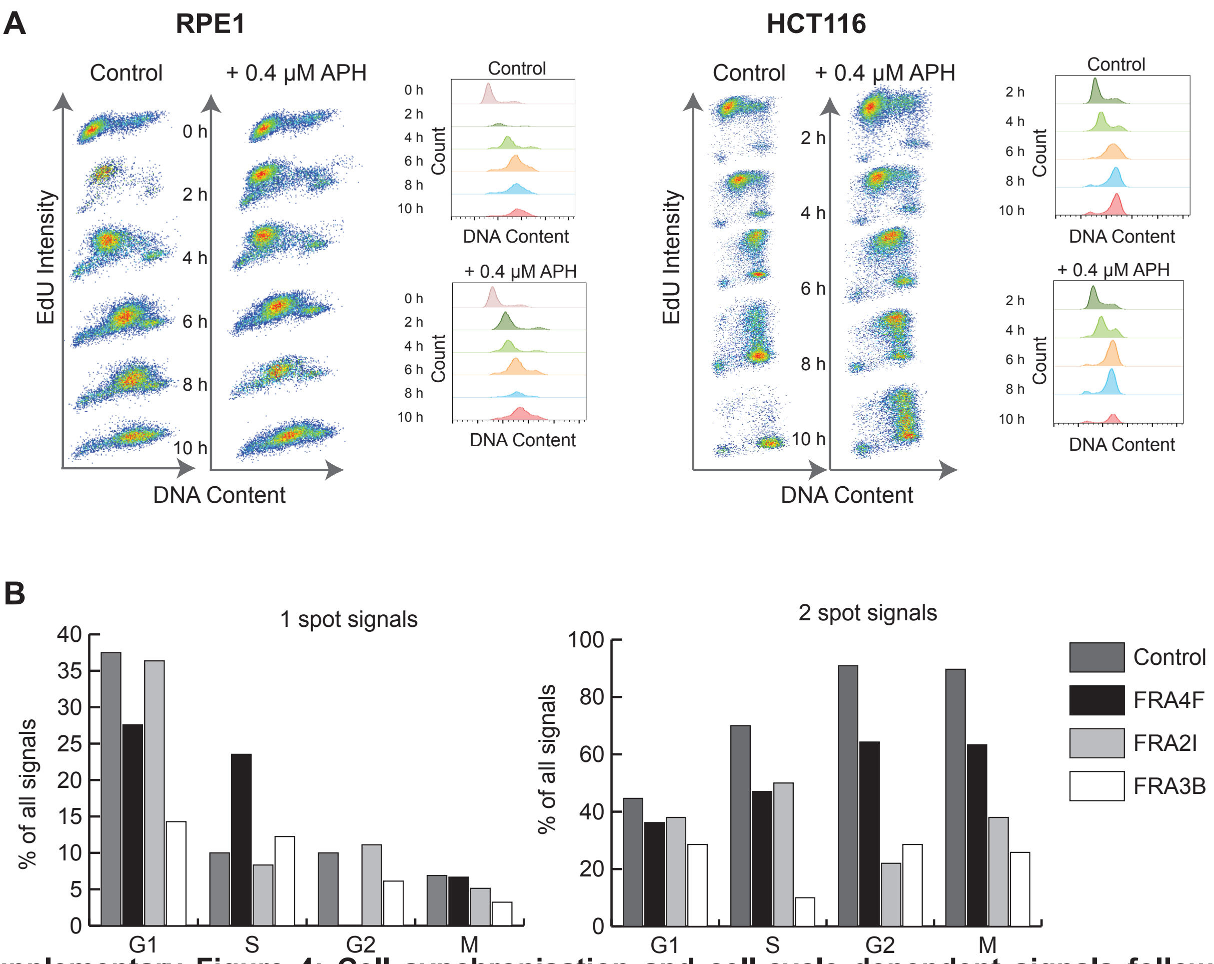


**Supplementary Figure 4: Cell synchronisation and cell-cycle dependent signals following premature chromosome condensation**

**A**. Cell cycle analysis, using flow cytometry, at different time points after release from G1/S block, as described in Figure 4A. Samples were pulsed with EdU 30 min prior to harvesting to mark replicating cells. EdU intensity versus DNA content (left) and propidium iodide (PI) histograms of the cell populations (right) are shown for different time points.

**B.** Frequencies of one-spot and two-spot signals at CFS locations and a control, non- fragile location, across different cell cycle stages in prematurely compacted chromosomes, using calyculin.

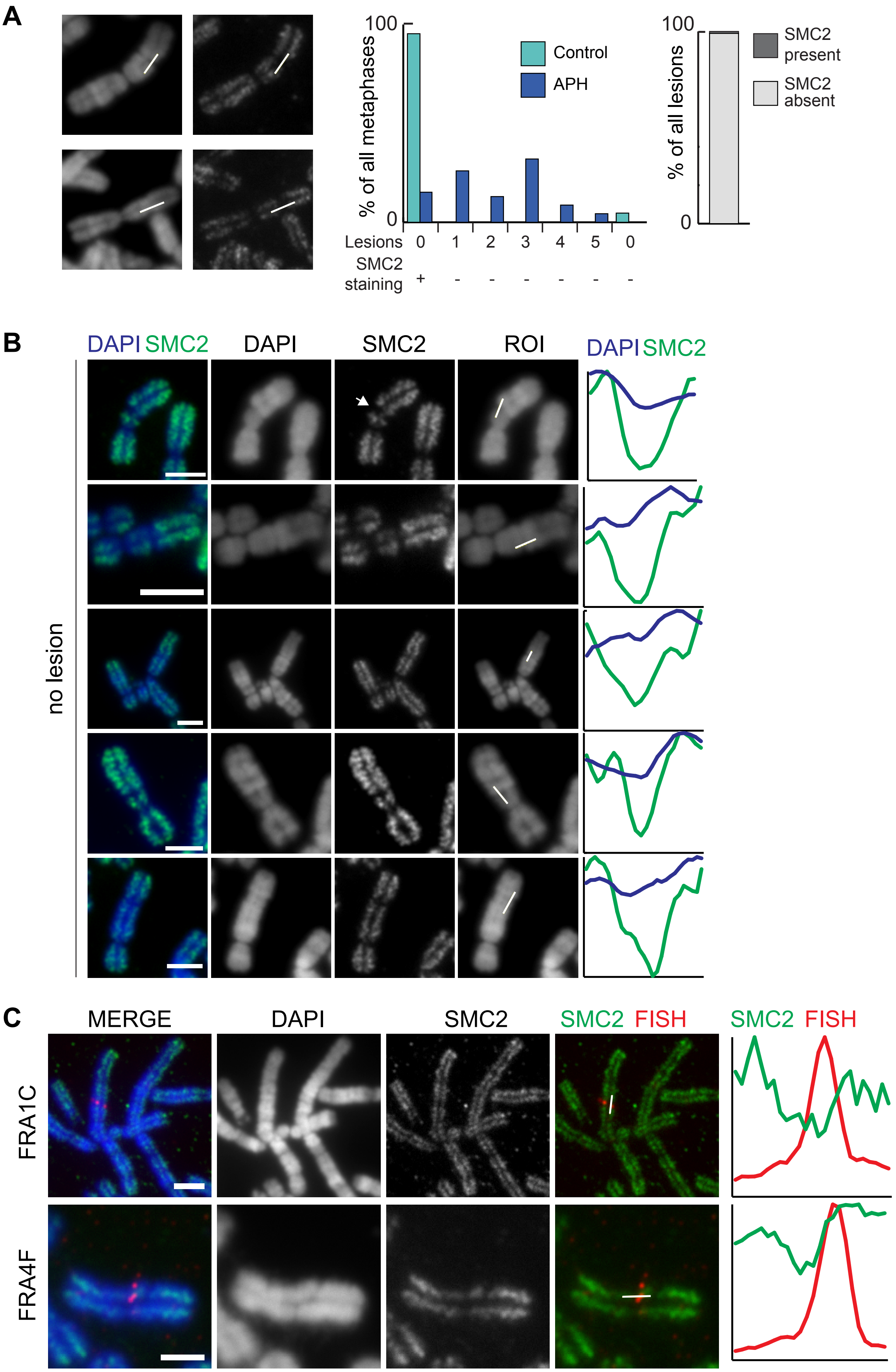


Supplementary Figure 5

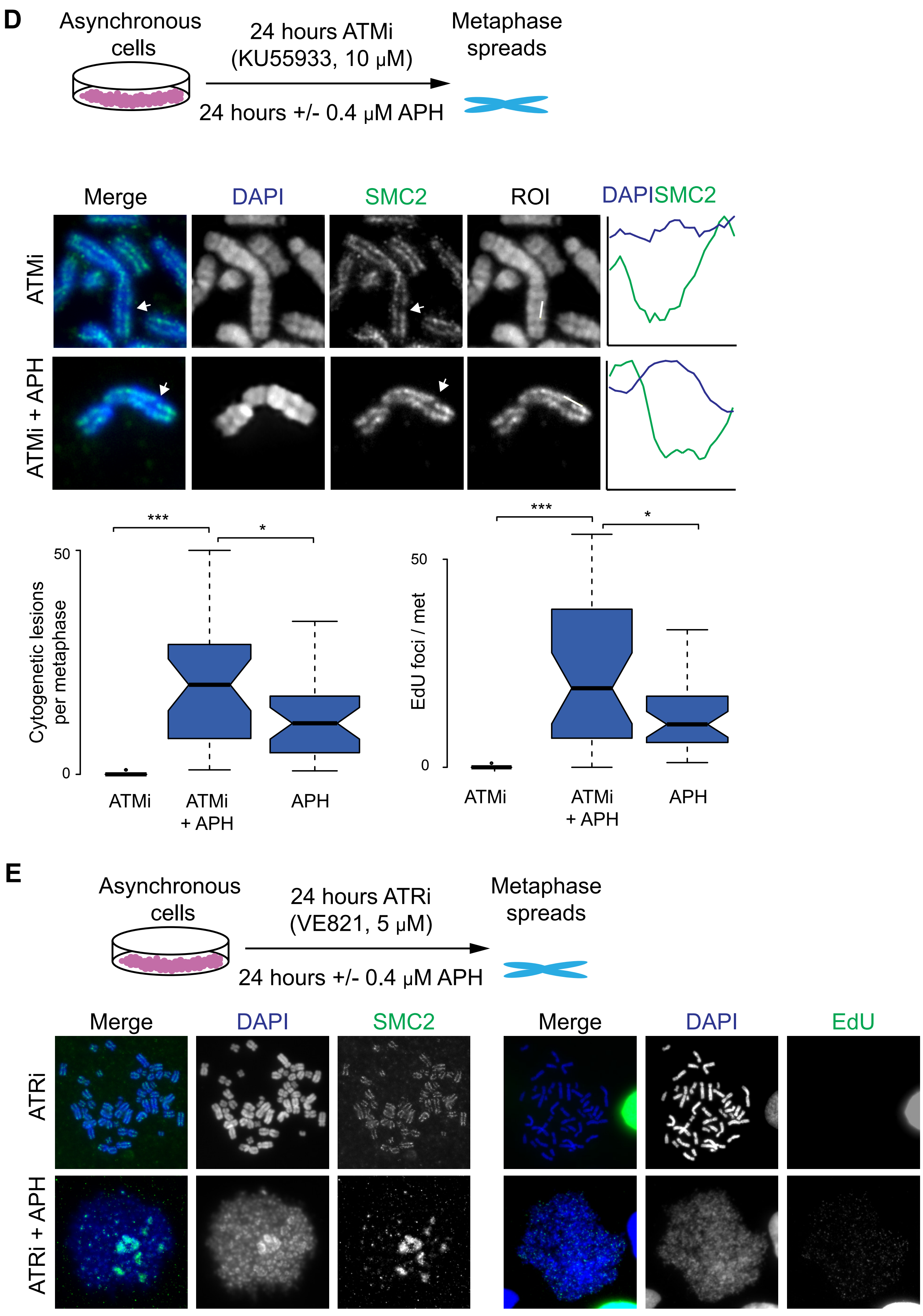


Supplementary Figure 5

**Supplementary Figure 5: SMC2 depletion at CFS loci on metaphase chromosomes.**

**A.** Regions of interest encompassing regions depleted of SMC2 used for intensity measurements in Figure 5A. Right, frequency of cytogenetic lesions and lesion-free SMC2 depletion (last column) in the presence or absence of aphidicolin in HCT116 cells. Far right, quantification of SMC2 occupancy at cytogenetic lesions in HCT116 cells.

**B.** Representative images and quantification of SMC2 depletion at cytogenetically normal chromosomes in RPE1 cells after aphidicolin treatment. Scale bar, 10 μm.

**C.** Representative immuno-FISH image showing a FISH probe for the FRA1C or FRA4F locus overlap with regions of SMC2 depletion on metaphase chromosomes from RPE1 cells or HCT116 cells; right, intensity profiles across the region of interest indicated by white line. Scale bar, 10 μm.

**D**. Top, diagram depicting experimental procedure to analyse effect of ATM inhibition on chromosome structure. Middle, representative images of cytogenetically normal chromosomes showing regions of SMC2 depletion following treatment with ATM inhibitor, and ATM inhibitor + aphidicolin; bottom, quantification of the number of cytological lesions (left) and MIDAS foci (right) in HCT116 cells following treatment with ATM inhibitor, aphidicolin and ATM inhibitor + aphidicolin.

**E**. Top, diagram depicting experimental procedure to analyse effect of ATR inhibition on chromosome structure. Bottom, representative images of chromosomes and fragmented metaphases in HCT116 cells following treatment with ATR inhibitor and ATR inhibitor + aphidicolin.

**F**. Boxplots showing normalised condensin binding in 1 kb windows within the FRA3B and 4q32.2 genomic loci at different time points in the absence (green) or presence (blue) of aphidicolin.

**G**. Top, line graphs showing mean normalised condensin binding in 1 kb windows at the FRA4F and FRA3O sites at different time points in the absence (green) or presence (blue) of aphidicolin. Bottom, boxplots of normalised condensin binding in 1 kb windows within the FRAF and the FRA3O sites across the different time points in the absence (green) or presence (blue) of aphidicolin.

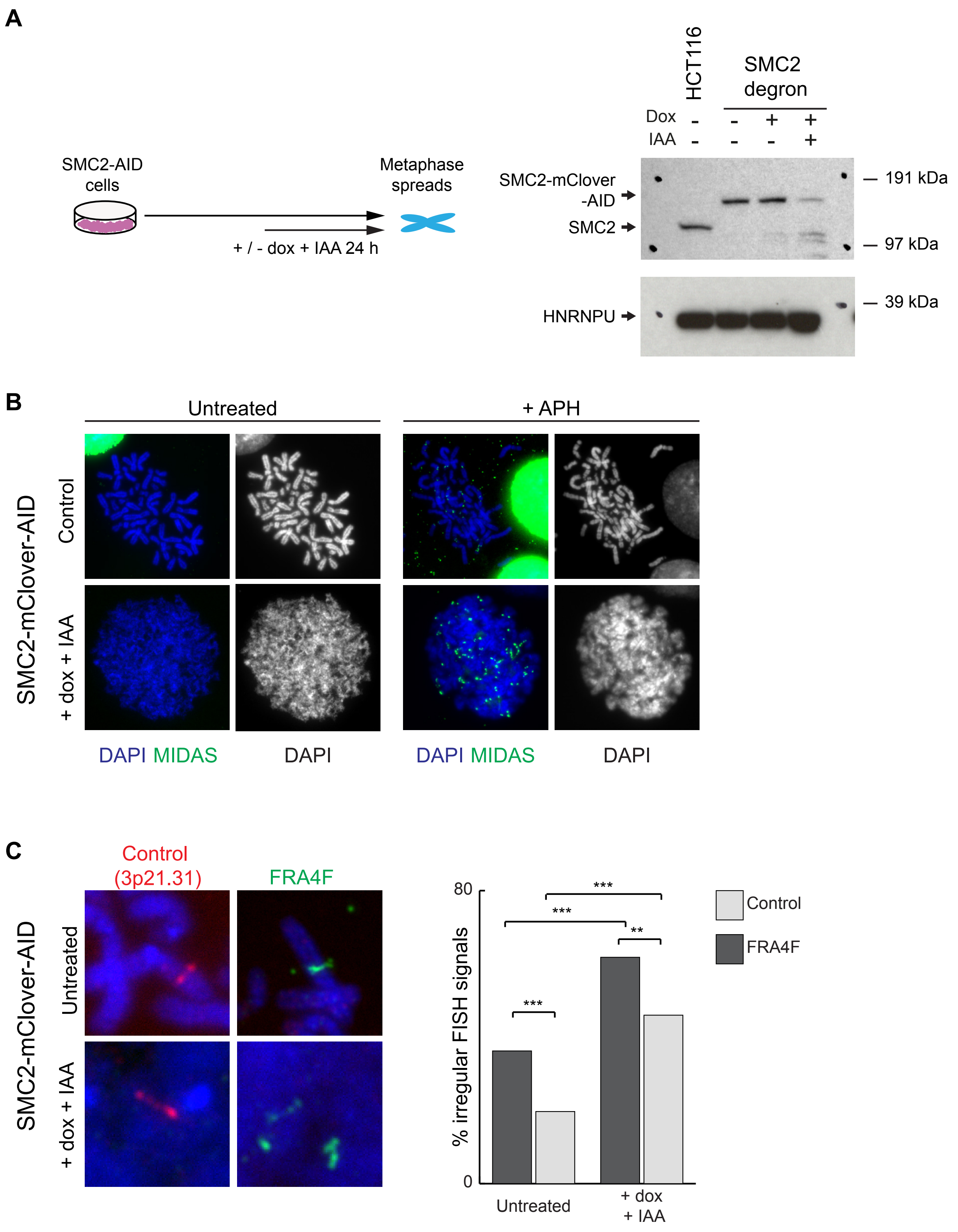


Supplementary Figure 6

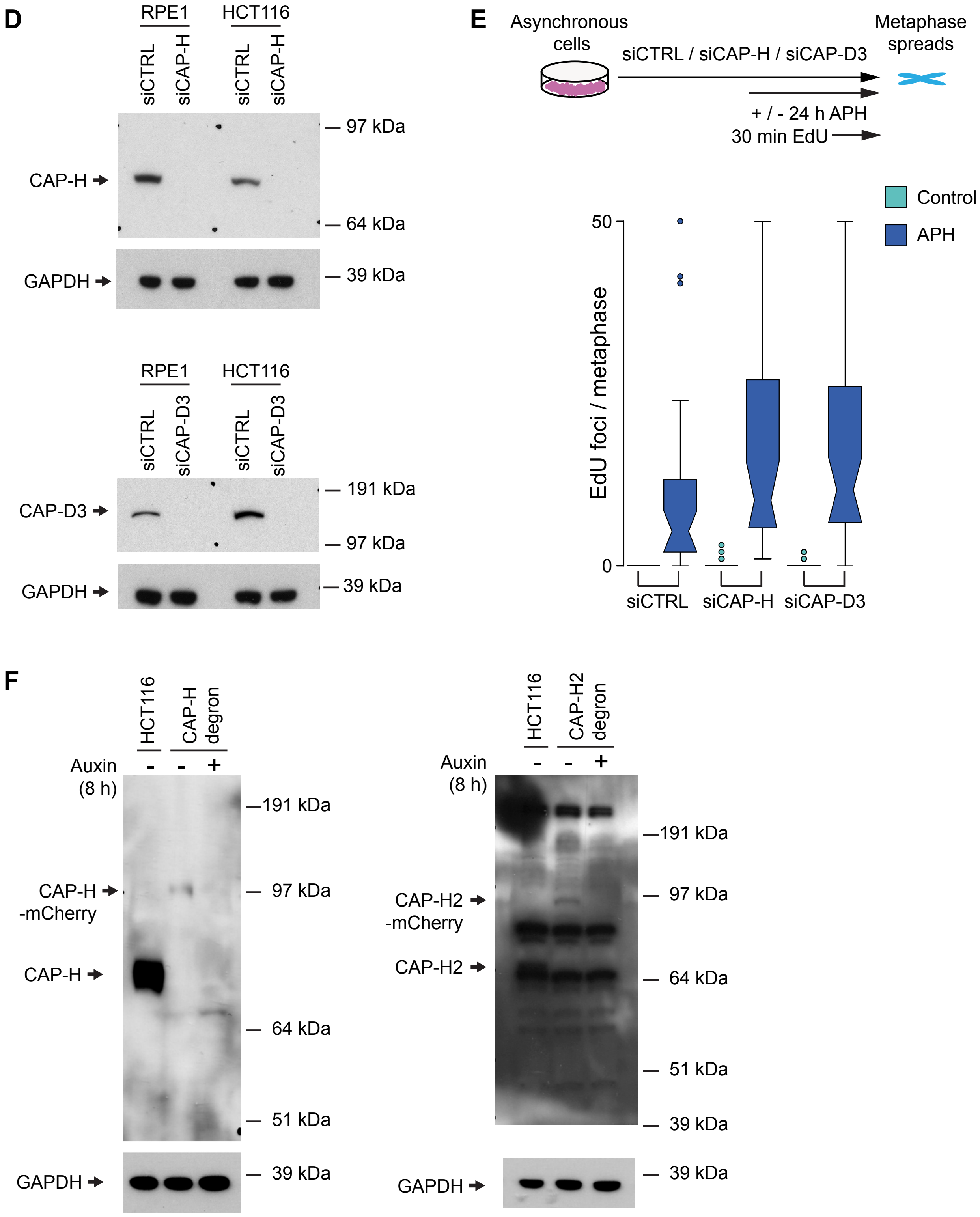


Supplementary Figure 6

**Supplementary Figure 6: Condensin depletion affects mitotic DNA synthesis and mitotic chromosome folding.**

**A**. Western blot showing depletion of CAP-H and CAP-D3 following 48 hours of siRNA treatment.

**B.** Boxplot showing the frequency of MIDAS foci per metaphase in HCT116 cells following CAP-H and CAP-D3 depletion in the absence (green) or presence (blue) of aphidicolin.

**C.** Top, diagram depicting auxin-induced SMC2 degradation in HCT116-SMC2-AID cell line. Bottom left, Western blot showing SMC2 degradation after auxin treatment. Bottom right, metaphase chromosome morphology and MIDAS labelling in the presence and absence of aphidicolin before and after auxin-induced SMC2 degradation.

**D**. Left, representative images showing FISH signals at the FRA4F fragile site and a control, non-fragile location in the HCT116-SMC2-AID cell line before and after auxin-induced SMC2 degradation. Right, quantification of the frequency of abnormal FISH signals before and after SMC2 degradation.
